# An algicidal bacterium shapes the microbiome during outdoor diatom cultivation collapse

**DOI:** 10.1101/2024.06.11.598583

**Authors:** Naomi E. Gilbert, Jeffery A. Kimbrel, Ty J. Samo, Anthony J. Siccardi, Rhona K. Stuart, Xavier Mayali

## Abstract

Biogeochemistry and productivity of algae-dominated environments is fundamentally influenced by the diversity and activity of bacteria. Namely, algicidal bacteria that prey on algal hosts can control elemental cycling and host populations within outdoor algal ponds used for biofuel production. In this study, we describe the genomic and proteomic signatures of a putative algicidal bacterium, *Kordia* sp. (family *Flavobacteriaceae*), that bloomed during a population-wide crash of the biofuel diatom, *Phaeodactylum tricornutum*. This *Kordia* sp. bloom occurred after 29 days of cultivation in outdoor algal raceway ponds inoculated with *P. tricornutum*, but not in parallel ponds inoculated with *Microchloropsis salina*. Several signatures of antagonism expressed by *Kordia* during diatom demise highlights previously unexplored mechanisms that may aid in algicidal activity or bacterial competition, including the type VI secretion system and hydrogen peroxide production. Analysis of accompanying downstream microbiota (primarily of the *Rhodobacteraceae* family) provides evidence that cross-feeding is important in supporting microbial diversity during algal demise. Specifically, *in situ* and laboratory data suggest that *Kordia* acts as a primary degrader of biopolymers during algal demise, and co-occurring *Rhodobacteraceae* exploit degradation molecules or scavenge metabolic byproducts for carbon. Further, targeted analysis of 30 *Rhodobacteraceae* metagenome assembled genomes suggest algal pond *Rhodobacteraceae* commonly harbor pathways for carbon monoxide oxidation, a potential strategy to persist under competition. Collectively, these observations further constrain the role of algicidal bacteria in the aquatic ecosystem.

## Introduction

Algal-bacterial interactions fundamentally influence aquatic biogeochemistry and ecology (1–4). Algae support a diverse bacterial community (microbiome) by exuding dissolved organic carbon (DOC) and bacteria support algae through metabolite production and nutrient remineralization, for example (5–11). Bacteria can also negatively impact algal hosts through competition, or directly *via* algicidal mechanisms (10, 12, 13). A better understanding of the feedback resulting from algal-bacterial interactions during algal growth and demise is needed to predict the role of the algal microbiome in natural and engineered ecosystems. Moreover, uncovering how bacteria interact with each other in the algal microbiome will allow more accurate quantification of resource recycling across the algal lifecycle.

Algal microbiomes are functionally diverse, leading to partitioning of DOC processing and assimilation (14–16). Furthermore, a succession in different bacterial taxa is well documented within marine algal blooms (17–20). Generally, in early algal growth, the DOC pool is composed of low molecular weight molecules (*e.g.,* amino/organic acids, sugars) exuded by the algae (2, 14, 21). This, in turn, enriches for small-molecule-users such as the *Rhodobacteraceae*, a group commonly found in algal microbiomes (14, 22). During algal demise, high molecular weight molecules (*e.g.*, polysaccharides, proteins, lipids) from lysis or exudation dominate the DOC pool (14, 23) and are directly consumed by biopolymer degraders, like the *Flavobacteriaceae* (19, 22, 24). Thus, a predictable succession of bacteria along the algal lifecycle is commonly observed.

Heterotrophic bacteria can influence the success of algal cultivation used for bioenergy production (10). Outdoor raceway ponds used to cultivate biomass are typically open-air and susceptible to the surrounding environment (25). Consequently, low pond productivity can result from suboptimal weather conditions and/or biological-driven demise, posing economic challenges (25, 26). Determining factors leading to low cultivation success and algal population crashes are of interest. Algal pond crashes can be driven by microbial pathogens (*e.g.*, viruses, algicidal bacteria, eukaryotic parasites) and/or environmental factors that have differing downstream effects on the biogeochemical and microbial fate of algal ponds (26, 27). Despite the need to identify the taxonomy, epidemiology and life history of pathogens causing algal demise, identifying the agent(s) that cause pond crashes remains challenging (27).

Here, we document the microbial underpinnings associated with an algal population crash that occurred during an outdoor cultivation effort. We followed a month-long time course of algal raceway (ARW) ponds inoculated with two distinct host algae: *Phaeodactylum tricornutum* (a diatom) and *Microchloropsis salina* (a chlorophyte). Using molecular tools (amplicon sequencing, metagenomics and proteomics), we characterized the microbiome dynamics across time for the two algal hosts. A crash was observed only in *P. tricornutum* ponds after ∼29 days of cultivation, which corresponded to a bloom in a single amplicon sequence variant assigned to the genus *Kordia* (family *Flavobacteriaceae*). We suspected that this bacterium contributed to *P. tricornutum* demise, as the *Kordia* genus contains several algicidal strains (28). We then profiled the microbiota following algal demise, with the initial hypothesis that microbial taxonomic and functional composition would mirror DOC availability along the algal life cycle. A laboratory experiment was performed to help explain patterns observed in the community-wide response to algal demise, suggesting cross-feeding is important in structuring the microbial community when algae die, specifically between marine *Flavobacteriaceae* and *Rhodobacteraceae*. Collectively, these results provide insight into *Kordia* sp. ecophysiology and emphasizes the importance of considering the role of bacteria-bacteria interactions within the algal microbiome, both in nature and industrial settings.

## Methods

### ARW pond time series design

Eight 557 L algal raceway ponds (ARW ponds) were inoculated with either *M. salina* (CCMP 1776, formerly *Nannochloropsis salina*; ARW4, 6, 9, & 11) or *P. tricornutum* (“Flour Bluff” isolate; ARW1, 5, 7, & 12) using natural, diatomaceous earth filtered seawater from Laguna Madre, Corpus Christi, TX on 10/27/2015 (Supplementary Figure 1). Cultures were amended with 2.0 mM NH_4_Cl, 2.0 mM pH balanced H_3_PO_4_, and 0.07 mM FeSO_4_. The first harvest (11/06/15) removed 75% of the cultures with media replacement and was followed by 50% harvests (with media replacement) on 11/10, 11/13, 11/17, 11/20, and 11/24. Samples for ash-free dry weight (AFDW) and biomass (for sequencing) were collected on 10/27, 11/2, 11/11, 11/17, 11/24, 11/27, and 12/2 (Supplementary Figure 1). Ph, salinity, temperature, and weather station data were collected twice daily.

### 16S/18S amplicons

Water samples, shipped overnight to LLNL on blue ice and immediately filtered onto 0.22 mm pore size SUPOR filters were extracted for DNA using Qiagen DNEasy kit. The 16S and 18S rRNA genes were amplified using 16S V4 primers 515F (29) and 806R (30) and 18S rRNA V4 primers 565F and 908R (31). Amplicons were sequenced through the Joint Genome Institute’s Community Sequencing Program (JGI CSP). DADA2 v1.6.0 (32) was used for quality control, and removeBimeraDenovo() was used to remove chimeras. Silva v132 (33) was used to assign taxonomy to the 18S/16S amplicon sequence variants (ASVs). The 16S ASVs were further assigned with RDP classifier v2.11 (34) against training set 16.

The R software platform (35) was used for statistics and data visualization [ggplot2 (36)]. Differential abundance of ASVs comparing day 1 *versus* day 29 was done with ANCOM-BC (v.2.2.2)(37). The VEGAN [v.2.6-4; (38)] package was used for the following: nonmetric multidimensional scaling (NMDS) based off bray-curtis distances, adonis() for significance of clusters, and diversity() for Shannon and Simpson’s indices. See Supplementary Methods for detail.

### Metagenomes

Total DNA from two *P. tricornutum* ponds (ARW1 and ARW7) sampled each timepoint was submitted for metagenomes with JGI’s CSP (Supplementary Figure 1). Raw reads were quality controlled using the JGI Standard Operating Procedure (39). Two co-assemblies were generated using QC’d reads across each pond using metaSPAdes v3.13.0 with kmers 21, 33 and 55 (40). Binning was done inside metawrap v1.3 (41) with metabat v2.12.1, maxbin v2.2.6, and concoct v1.0.0. Bins were assessed with checkM v1.1.3 (42), and those with a completion > 50% and contamination <10% were kept and reassembled as individual assemblies using SPAdes v3.13.0 (43). Resultant metagenome assembled genomes (MAGs) were dereplicated into a set of non-redundant MAGs using dRep v2.2.3 (44). To estimate MAG relative abundances, QC’d reads were mapped to the MAGs with bbmap v35.85 (45) and perfectmode=t. MAG abundance was calculated using the mean of the median fold coverage (“MAG abundance”) in each sample (46).

MAG taxonomy was assigned with GTDB-tk v.1.0.0 r202 using default parameters (47). The PATRIC pipeline (48, 49) was used for gene calling and functional annotation. GATOR (https://github.com/jeffkimbrel/gator) was used in parallel to assess metabolic pathway completeness, curated for pathways relevant to algal microbiomes and unique carbon/energy utilization pathways. Carbohydrate-active enzyme (CAZyme) HMMs were queried using dbcan12 with dbCAN2 (50). Orthofinder 2.5.5 (51–53) was used to find orthogroups and a species tree was generated using default parameters and annotated using iTol v5 (54).

### Metaproteomes

Total proteins were extracted following Mayali *et al.* 2023 (11). A Waters nano-Acquity dual pumping UPLC system (Milford, MA) was configured for on-line trapping of a 5 µL injection at 5 µL/min with reverse-flow elution onto the analytical column at 300 nL/min. MS analysis was performed using a Q-Exactive HF mass spectrometer (Thermo Scientific, San Jose, CA) outfitted with a home-made nano-electrospray ionization interface. See Supplemental methods for more detail. Protein spectra were mapped to a database of assembled MAGs (1,134,297 sequences), a *P. tricornutum* genome [10,408 sequences, *Phaeodactylum tricornutum* CCAP1055.1 (55)], public *M. salina*/*Nannochloropsis* genomes (10, 964 sequences, NCBI TaxID 5748) obtained March 2021, and 16 common contaminant proteins. Sequences were processed using Protein Digestion Simulator (https://github.com/PNNL-Comp-Mass-Spec/Protein-Digestion-Simulator). MSGF+ (56) was used to identify peptides against the custom protein database in target/decoy mode with 20 ppm parent ion tolerance, tryptic rule without post-translational modifications considered. Best MSGF+ search matches were filtered at 1% FDR and MASIC (57) was used to pull identified peptide abundances. Only protein specific peptides (peptides unique to protein in the whole protein collection) were used in consequent analysis and aggregation. Protein abundance is reported as normalized spectral abundance factor (NSAF) as described in (58).

The *Kordia* sp. (mARW1_16) expressed proteins were manually assigned KEGG functional categories using the PATRIC annotations. GATOR pathways were merged with this set of proteins and data were plotted using ggplot2 (36).

### Cross-feeding laboratory experiment

We tested whether pre-conditioning of *P. tricornutum* lysate (analog to demise conditions) by *Kordia algicida* OT1 (OT1, primary degrader) enhances growth of a representative *Rhodobacteraceae* strain (*Sulfitobacter* sp. N5S). OT1 was purchased from the Japanese National Institute of Technology and Evaluation (NITE) Biological Resource Center (NBRC Accession #100033). We annotated the public genome as described above. N5S was isolated from Bodega Bay, CA (38.332615, −123.048296) by spreading whole seawater onto Marine Agar plates. See the Supplemental Methods for detail.

*P. tricornutum* CCMP 2561 lysate was generated using exponentially growing culture in sterile F/2 media as described in Supplemental Methods. Algal cells were pelleted by spinning at 5000 rpm for 8 min and the spent media discarded to remove algal exudate. Cells resuspended in sterile F/2 were lysed using an Ultrasonic Processer XL sonicator (Misonix) and filtered through a 0.8 µm syringe filter. The final lysate was then incubated with or without OT1 to produce “abiotic” and OT1 “conditioned” lysate. The conditioning experiment was done in triplicate in 3 mL sterile borosilicate tubes at 22°C in the dark for 5 days. Growth was monitored using OD600 (BioTek Cytation 5 plate reader, Aglient) and flow cytometry. OT1 and N5S was then inoculated into 0.2 µm filtered abiotic *versus* conditioned lysate in 5 replicates in a 96 well plate (rounded bottom). Negative controls per condition and a positive control (Zobell) was run. OD600 was read every 20 min for 72 h using a BioTek Cytation 5 plate reader at 22°C in the dark. Flow cytometry samples were collected at T=0 h and T=72 h. The t-test was used to compare cell abundances at T= 72h using GraphPad Prism 9. See Supplemental Methods for more detail.

## Data Availability

DNA/amplicon sequencing data are available through the JGI’s Integrated Microbial Genome database under accession codes in Supplementary Table 2. MAG assemblies are available *via* Zenodo (https://doi.org/10.5281/zenodo.11414289). Bacterial strains are available upon request. Raw proteomics data are available on MassIVE (accession# MSV000094933).

## Results

### Differential persistence of the two algal species and evidence of P. tricornutum population decline in the ARW ponds

During concurrent cultivation of 8 co-located ARW ponds (4 inoculated with the diatom *Phaeodactylum tricornutum*, and 4 inoculated with green alga *Microchloropsis salina*), an algal crash occurred only in the four *P. tricornutum* ponds after 27 days of semi-continuous cultivation (that included 6 harvests of biomass with media replenishment). A visible change in color from dark brown (healthy *P. tricornutum* biomass) to a milky white was noted. A steady decline in algal biomass (∼0.3 to <0.1 g/L) across *P. tricornutum* ponds occurred (Supplementary Figure 2, Supplementary Table 3), but not for *M. salina* after day 29 (Supplementary Figure 2). We used 18S rRNA gene profiling, focusing on the genera assignments *Microchloropsis* or *Phaeodactylum* within their respective ponds. *Microchloropsis* ponds exhibited stable host 18S rRNA relative abundance, whereas *P. tricornutum* ponds significantly declined in the relative abundance of host 18S rRNA at day 33 (5.8-log fold change between day 1 *versus* day 33 [ANCOMBC, p_adj_=0.0001]; Figure 1A & Supplementary Table 3). Proteomics data indicate a declining trend in proteins identified as *P. tricornutum* on days 29 and 33, with negligible proteins identified as *M. salina* (Supplementary Figure 3). Therefore, we refer to this event as a population-wide “crash”, with “demise” referring to the steady decline in AFDW of *P. tricornutum* after day 16 (Supplementary Figure 2).

**Figure 1.**
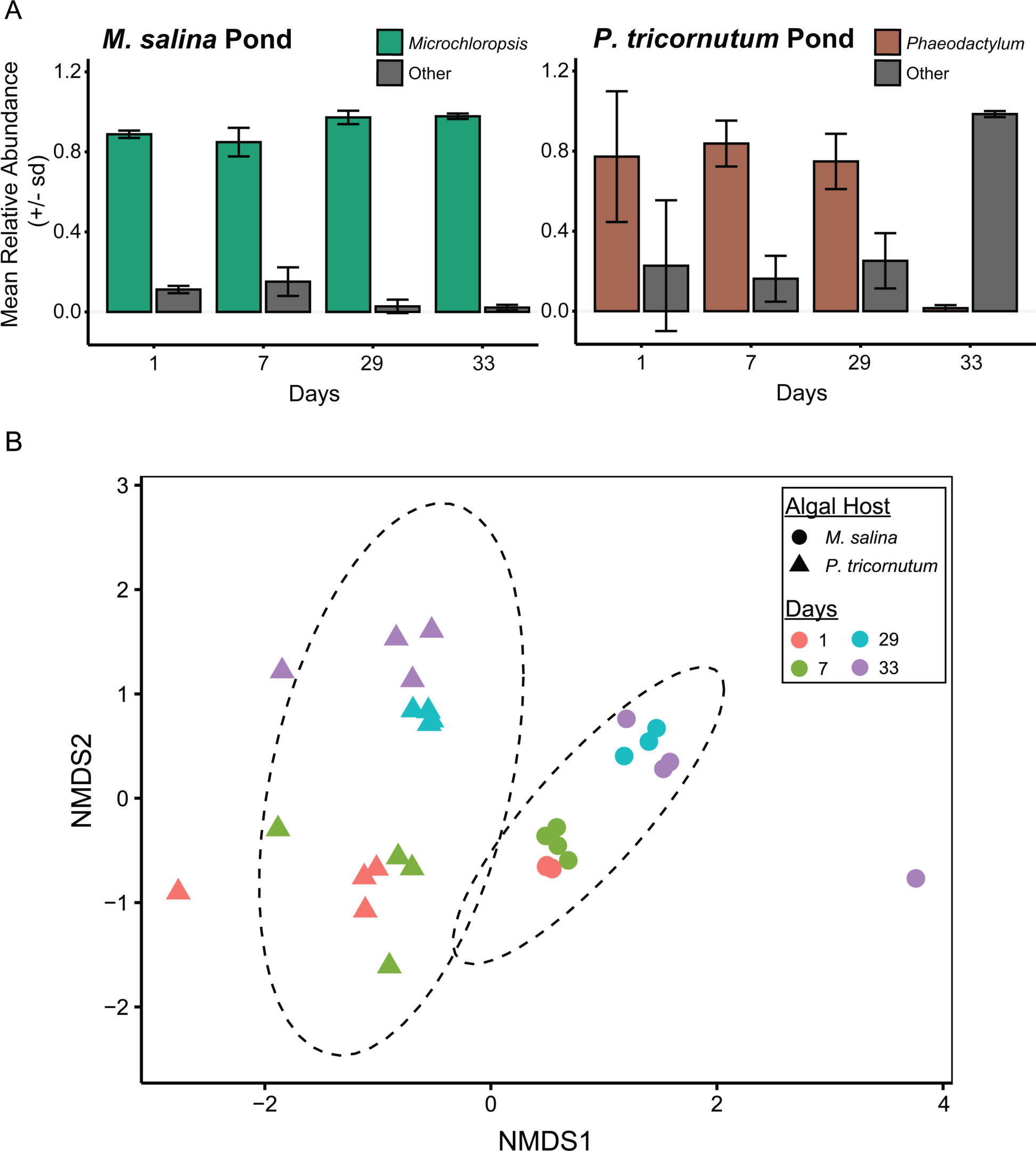
Algal host strain and decline in *P. tricornutum* over time influence microbiome composition within the ARW ponds. A) Relative abundance of representative host 18S rRNA ASVs between ponds inoculated with *M. salina versus P. tricornutum*. Error bars represent the standard deviation in relative abundance between four replicate ponds. 18S rRNA amplicons assigned to either *Phaeodactylum* or *Microchloropsis* are shown in either brown or green, where all other 18S rRNA amplicons are shown as “Other” in gray. B) NMDS ordination based on the Bray-Curtis dissimilarity matrix of 16S rRNA bacterial communities between samples inoculated with *M. salina* (circles) and *P. tricornutum* (triangles) over timepoint (indicated by color); 95% confidence ellipses (stat_ellipse, Fox and Weisberg 2011) are grouped by algal host inoculum. Stress = 0.115.

### Bacterial community composition responded to algal host species and health

Algal host ponds had distinct microbiome compositions that shifted with time (Figure 1B), especially pronounced for *P. tricornutum* ponds (Supplementary Figure 4; presumably due to their collapse). Non-metric Multi-dimensional Scaling (NMDS) ordination of the Bray-Curtis dissimilarities clustered 16S rRNA gene microbiome profiles based on algal host (Figure 1B, ADONIS p<0.0001). The dominant bacterial groups differed between host-alga (Supplementary Figure 5). *M. salina* ponds were dominated by ASVs assigned to the Alpha- and Betaproteobacteria, Sphingobacteriia, and “unclassified” Pseudomonadota (Supplementary Figure 5). The *P. tricornutum* ponds were dominated by ASVs assigned to the Alpha-, Gamma- and Betaproteobacteria at days 1 and 7, but shifted to predominantly Flavobacteriia and Alphaproteobacteria classes at days 29 and 33 during algal demise and population crash (Supplementary Figure 5).

We further inspected the microbiome of the *P. tricornutum* ponds due to evidence of algal population demise. Based on NMDS clustering of beta-diversity with time, two major clusters of ASVs in the *P. tricornutum* microbiome emerged (Supplementary Figure 4a; ADONIS p<0.005). These clusters correspond to *P. tricornutum* growth phases across the ARW ponds: “growth” (days 1 & 7) and “demise” (days 29 & 33). The alpha-diversity of the demise microbiome was also significantly reduced compared to the growth microbiome (Supplementary Figure 4B).

### Taxon-specific response to P. tricornutum demise – identification of a highly abundant potential pathogen, Kordia sp

Examination of individual ASVs across the *P. tricornutum* growth phases revealed an ASV-specific succession (Figure 2, Supplementary Figure 4A). One ASV uniquely assigned to the genus *Kordia* (ASV_4) dominated the demise phase (reaching up to 93% relative abundance of the 16S rRNA reads, Figure 2), and had a 12.5-log fold increase in abundance on day 33 *versus* day 1 (ANCOMBC, p_adj_ =3.47e-75, Supplementary Table 4). ASV_4 was nearly undetectable during growth timepoints (Figure 2) and had no detectable reads in the *M. salina* microbiome (Supplementary Figure 6). Aside from ASV_4, the demise microbiome was primarily composed of many ASVs assigned to the family *Rhodobacteraceae*, with ASV_24 reaching up to ∼6% relative abundance within a sample (Figure 2). On the contrary, the growth microbiome contained a greater diversity of family-level taxonomic affiliations, with non-*Kordia Flavobacteriaceae*, *Alteromonadaceae,* and *Sphingomonoadaceae* contributing high relative abundances (Figure 2). Several *Rhodobacteraceae*, *Stappiaceae* and *Methylophagaceae* - assigned ASVs persisted throughout both phases (Figure 2). In summary, the microbiome shifted from a diverse “healthy” microbiome during growth to one dominated by two main taxonomic groupings during demise and population crash: *Flavobacteriaceae* (*Kordia* sp.) and the *Rhodobacteraceae* (Figure 2).

**Figure 2.**
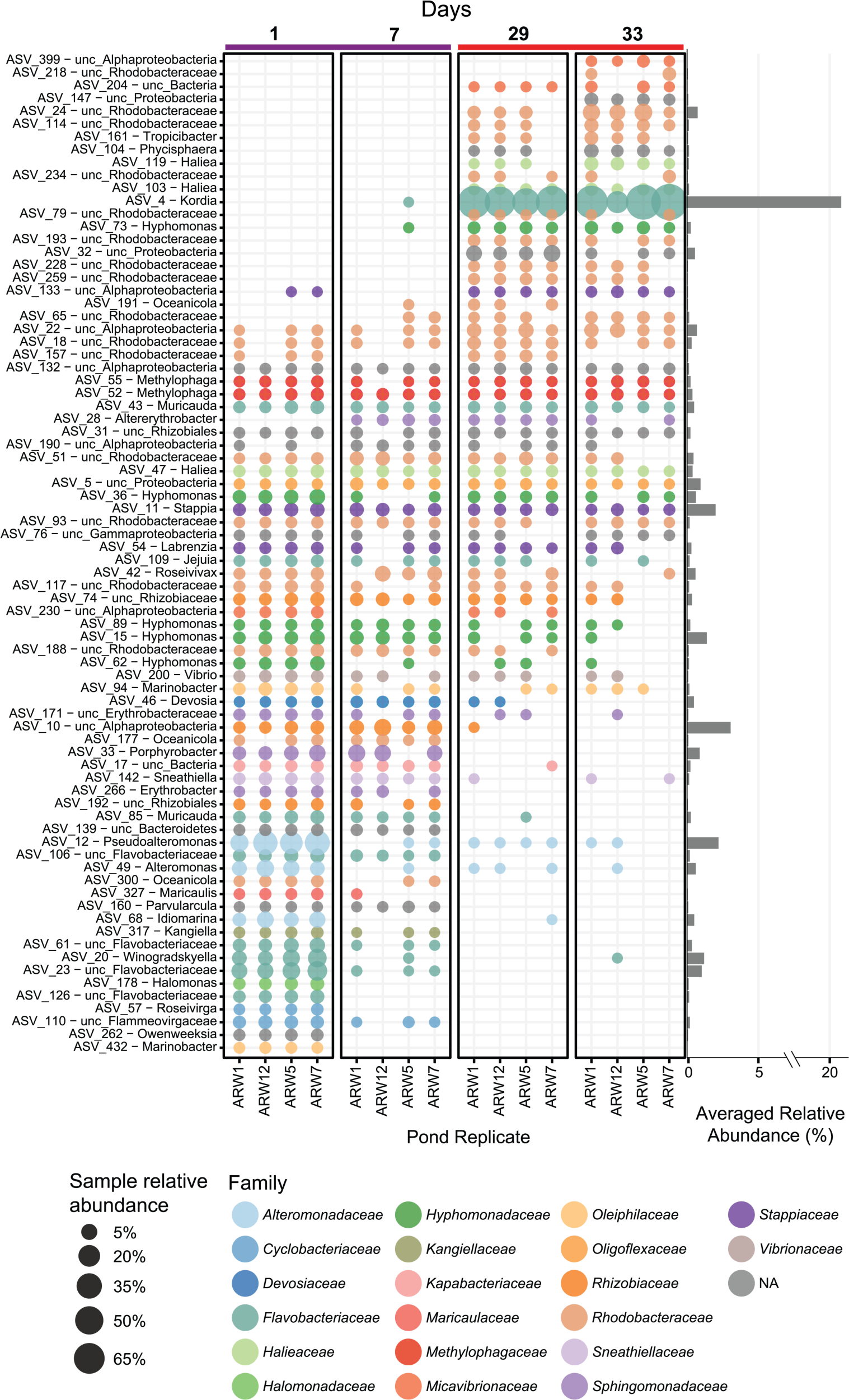
Shift in microbiome composition during *P. tricornutum* demise corresponds to a bloom of an individual ASV assigned to the genus, *Kordia*, and co-occurring *Rhodobacteraceae* taxa. Each pond replicate is shown on the x-axis and individual ASVs (shown with ASV number and lowest taxonomic assignment) are on the y-axis. The timepoints are grouped and color-coded by algal growth phase, which is based on NMDS clustering of the relative abundance of ASVs in Supplementary Figure 4 (growth [purple line] versus demise [orange line]). Each bubble represents the relative abundance value for each replicate, and the averaged relative abundance across all the plotted samples for each ASV is shown on the bar graph to the right. Bubbles are color coded by the taxonomic assignment to the Family level.

Metagenome-assembled-genomes (MAGs) were generated for 2 of 4 raceways from the *P. tricornutum* ponds (“ARW1” and “ARW7”, Supplementary Table 5). A total of 143 non-redundant MAGs were assembled, some of which cluster with bacterial strains previously isolated from these ponds and co-cultured with *P. tricornutum* [(11, 59); Figure 3]. Taxa of the phyla Myxococcota, Planctomycetes, Bdellovibrionota, Patescibacteria, Bacteroidota, Gammaproteobacteria and Alphaproteobacteria were also detected (Figure 3). Within the Alphaproteobacteria, there was high representation (33 MAGs) of the *Rhodobacteraceae* (Figure 3). As expected, the *Kordia* MAG (mARW1_16, Figure 3) had the highest MAG abundance values across the *P. tricornutum* ponds (average 28% relative abundance, Figure 3). Like the amplicon data, *Flavobacteriaceae* and *Rhodobacteraceae* family-level assignments were the dominant groups during the demise phase, whereas the growth phase had representation from more diverse taxa (Supplementary Figure 7). However, MAGs of the *Pirellulaceae, Rhizobiaceae,* and *Phycisphaeraceae* families were also present in the demise phase, although in lower abundances (Supplementary Figure 7).

**Figure 3.**
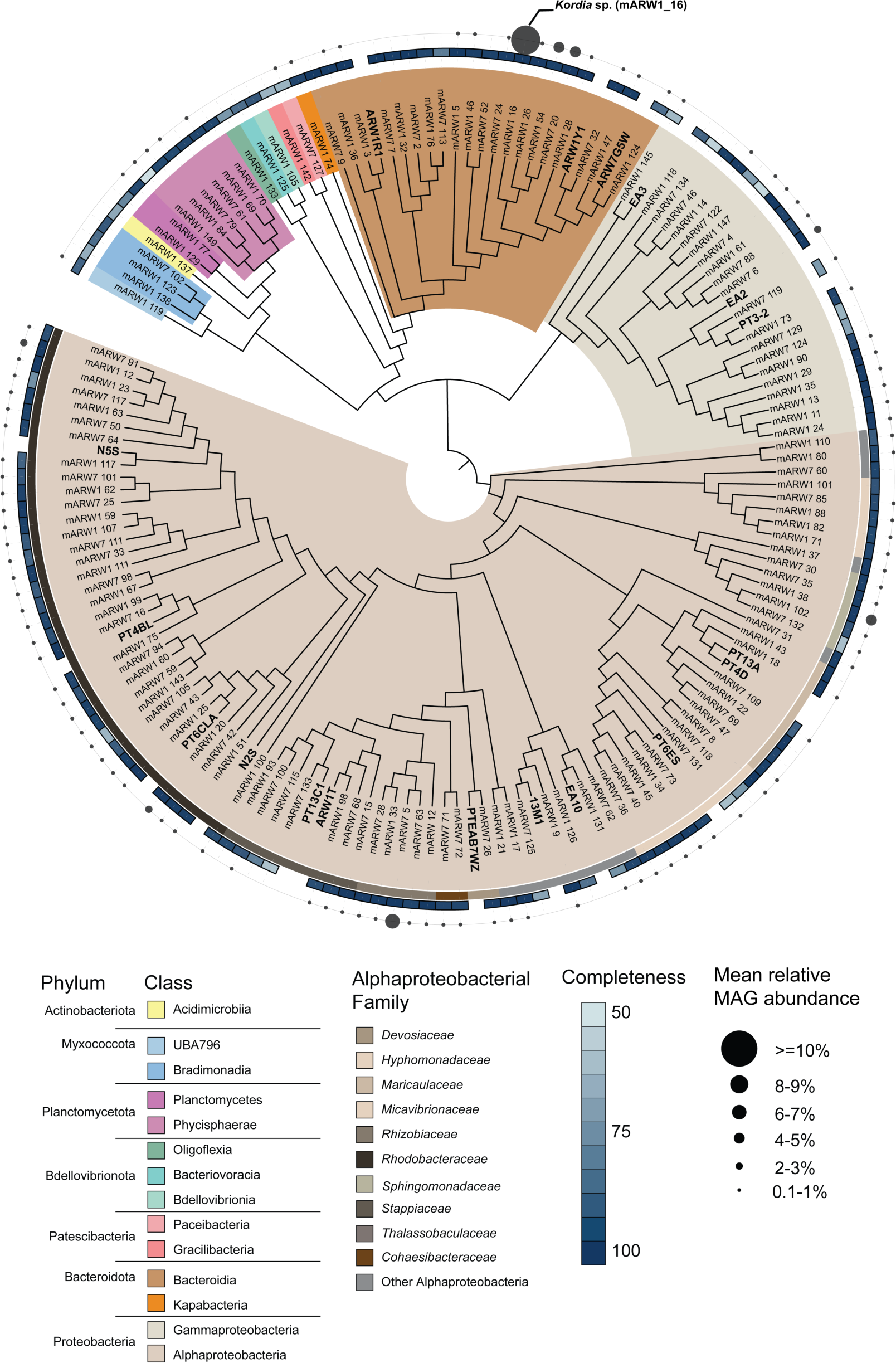
Distribution of MAGs detected in the *P. tricornutum* pond metagenomes. The phylogenetic tree shows the relationship between the MAGs based on orthogroups (see Methods). Cultivated isolates are denoted by their strain identification around the tree as follows: PT13A: *Oceanicaulis* PT13A, ARW1R1: *Algoriphagus* ARW1R1, ARW1Y1: *Muricauda* ARW1Y1, ARW7G5Y1: *Arenibacter* ARW7G5Y1, PTEAB7WZ*: Devosia* PTEAB7WZ, ARW1T: *Stappia* ARW1T, PT6CLA: *Rhodophyticola* PT6CLA, PT4BL: *Yoonia* PT4BL, EA3: *Pusillimonas* EA3, EA2: *Alcanivorax* EA2, PT6ES: *Henriciella* PT6ES, EA10: *Tepidicaulis* EA10, PT13C1: *Roseibium* PT13C1, 11-3: *Thalassospira* 11-3, PT3-2: *Marinobacter* 3-2, ARW7G5W: *Muricauda* ARW7G5W, PT4D: *Oceanicaulis* PT4D, N2S: *Roseobacter* N2S, N5S: *Sulfitobacter* N5S. The relative abundance range of MAGs averaged across all samples are shown as bubbles surrounding the tree. MAG completeness shown as a heatmap ranging from 50-100% completeness. Clades are color-coded by Class-level GTDB taxonomy, and to the Family-level for Alphaproteobacteria.

### Expressed proteins mapped to Kordia sp. provide insight into mechanism leading to its dominance during algal pond demise

Using metaproteomics of *P. tricornutum* pond samples over time, we identified expressed proteins which mapped to the *Kordia* MAG (mARW1_16) to gain insight into potential factors linking *Kordia* to diatom demise. There were 1,728 of 4,252 (40.64%) proteins in the mARW1_16 MAG detected in the metaproteome (Supplementary Figure 8, Supplementary Table 6). Proteins in the Transcription/Translation, Transport, Amino acid metabolism, and Central carbon metabolism categories were most detected, suggesting the *Kordia* population was metabolically active (Figure 4A). The next most abundant categories were the Peptidases/proteases & inhibitors and Complex carbon metabolism categories (Figure 4A). Further, the Complex carbon metabolism category harbors the most abundant protein detected (based on normalized spectral abundance factor, NSAF) across demise/crash samples: a SusC-family Ton-B Dependent Transporter (Figure 4B). The gene encoding this protein is located nearby genes encoding glycoside hydrolase family 2 TIM barrel-domain, beta-glucanase, and a putative lamarinase targeting beta-glucan, all detected in the *Kordia* proteome (Supplementary Table 7). Likewise, several proteins assigned as SusD surface glycan-binding PUL component (Complex carbon metabolism) were abundant (Figure 4B).

**Figure 4.**
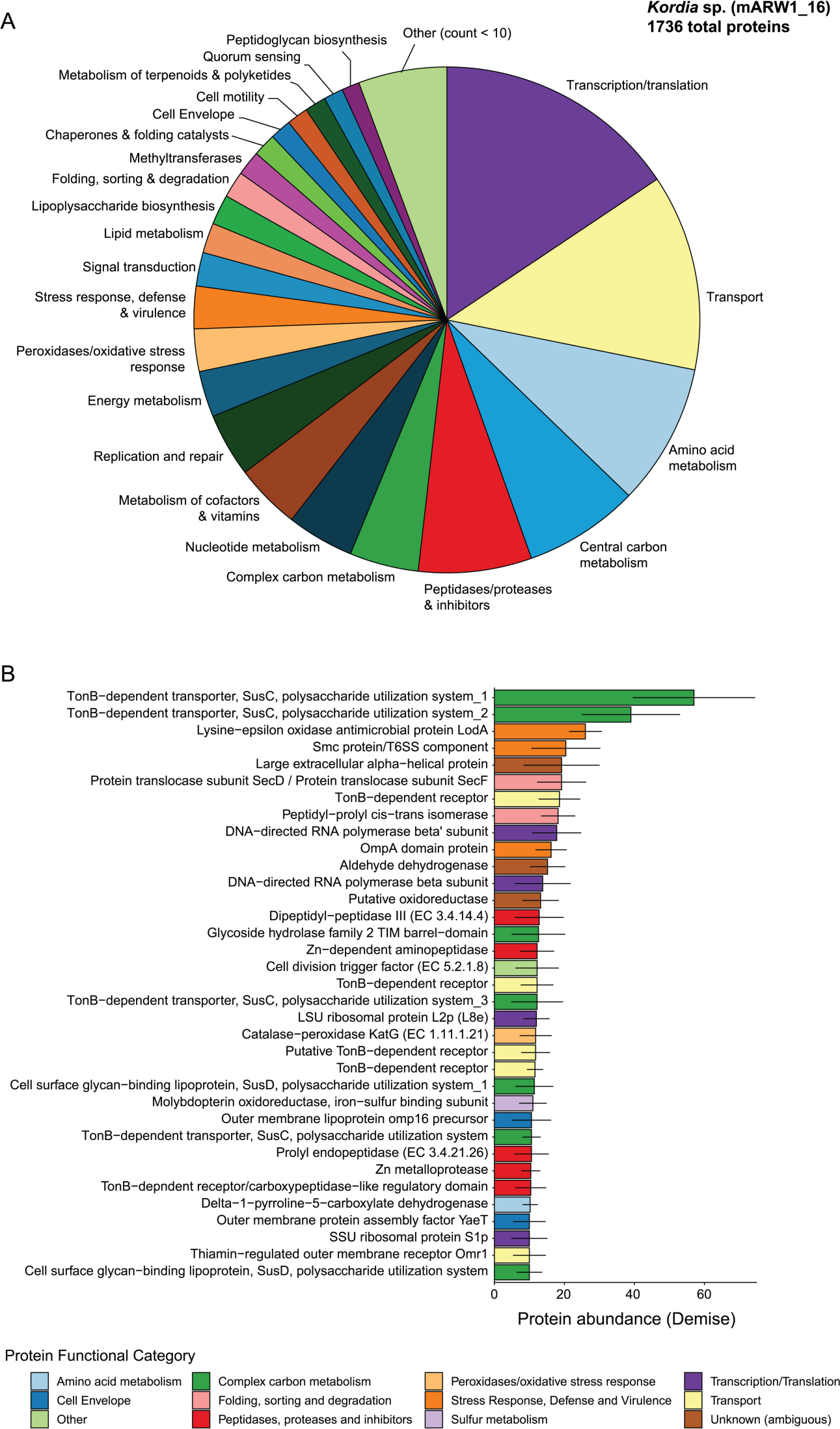
Abundant protein functional categories and proteins mapped to the *Kordia* MAG indicate antagonism and biopolymer degradation. A) Breakdown of the most detected protein functional categories mapped to Kordia. The “Other” category specifies categories with less than 10 detected proteins across all metaproteomics samples. Proteins with “unknown” or “ambigous” function were excluded for visual purposes. Protein assignments are available in Supplementary Table 6. B) The topmost abundant *Kordia* proteins detected across the demise samples (day 29 and 33) that had an average NSAF >= 10. Average NSAF across all demise samples (n=5) are shown and their standard deviations are represented as error bars.

The next two most abundant proteins belonged to stress response, defense and virulence (Figure 4B). The lysine-epsilon oxidase protein (LodA), known for its antimicrobial activity resulting from hydrogen peroxide production (60) was among the top three most abundant *Kordia* proteins expressed across algal demise (Figure 4B, Supplementary Table 6). Additionally, a putative type VI secretion system (type 6SS) component was the fourth most abundant protein (Figure 4B) and is located nearby two expressed type 6SS baseplate protein components (TssJ and VgrG) and many other encoded T6SS components (Supplementary Table 6).

### Genomic analysis indicates Kordia fills a specific niche compared to the co-occurring Rhodobacteraceae community

We initially hypothesized that microbial taxa present during the demise phase may have metabolized complex biomolecules expected to be released from dead or dying diatom cells. We then asked whether taxa in the demise microbiome, other than *Kordia* sp., were specialized to acquire carbon from biopolymers. We examined the CAZyme profiles of *P. tricornutum* pond MAGs, focusing on comparing the *Kordia* sp. MAG to all MAGs assigned to the *Rhodobacteraceae* family, as these two groups dominated the demise microbiome (Figure 2, Supplementary Figure 7). We counted all CAZyme hits to glycoside hydrolases (GH) and polysaccharide lyases (PL) and summarized them at the individual MAG level (Figure 5A). We found that *Kordia* sp. harbored ∼255 of these CAZymes, while the *Rhodobacteraceae* MAGs averaged ∼61 CAZymes per MAG (n=30 MAGs, Figure 5A). In the *Kordia* MAG specifically, CAZymes were predicted to target a diverse range of substrates such as beta-glucan (>10), chitin/peptidoglycan (>7), glycogen (5) and several others (Figure 5B). Some *Rhodobacteraceae* MAGs harbored a diverse repertoire of CAZymes (*e.g.,* mARW1_75, mARW1_60), however MAGs with specific genus-level matches to the ASVs grouped in the demise phase were relatively depleted in diverse CAZymes (Figure 5A). This suggests that although *Rhodobacteraceae* members prevailed under diatom demise, they did not universally harbor the capability to degrade diverse complex carbon sources.

**Figure 5.**
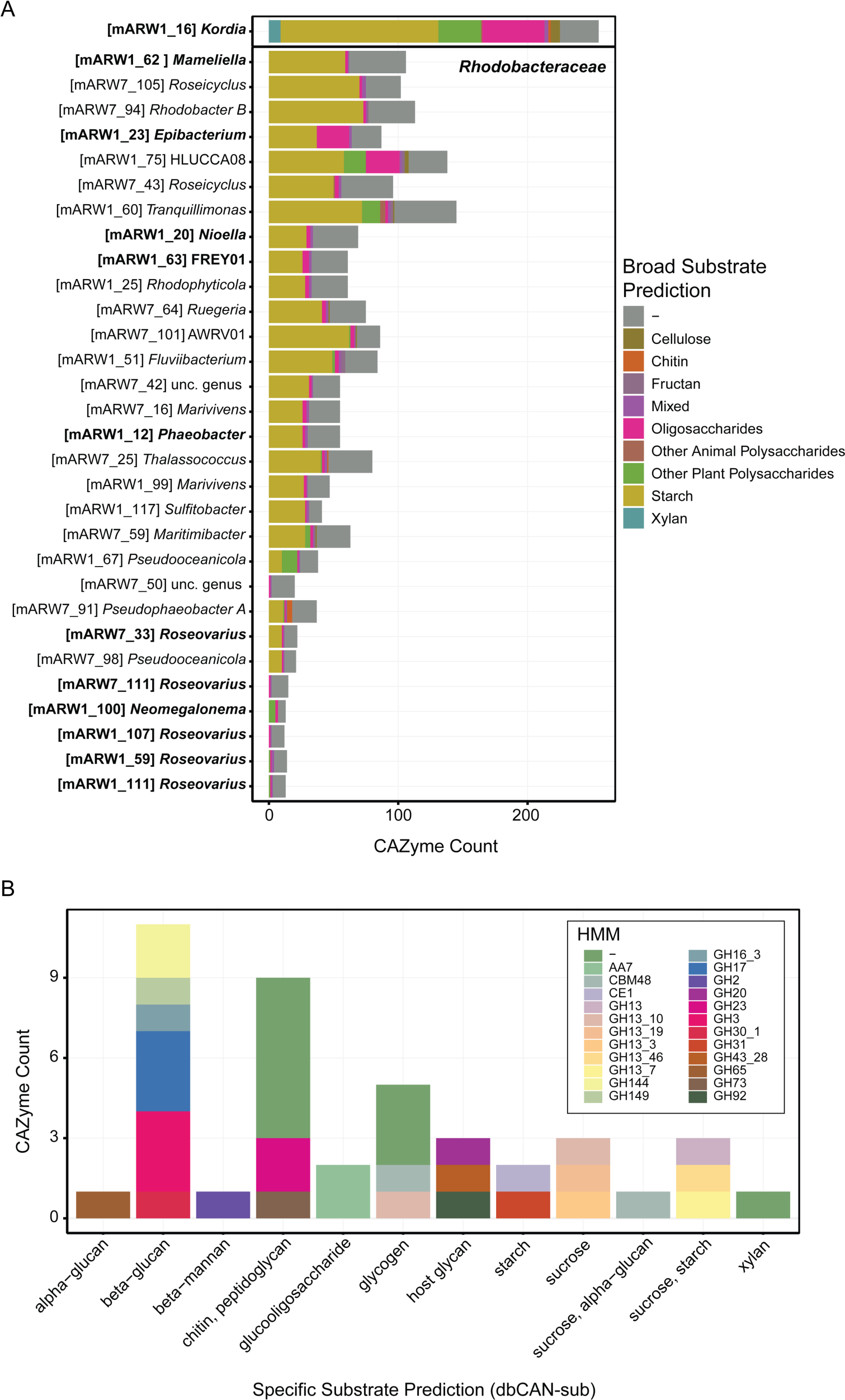
*Kordia* fills a distinct niche compared to *Rhodobacteraceae* during algal demise by encoding abundant CAZymes targeting a diverse range of predicted biopolymers. A) All *Rhodobacteraceae* non-redundant MAGs at >= 75% completion (n=30) assembled from either the ARW1 or ARW7 ponds are shown for comparison with the *Kordia* MAG (boldened). MAG IDs are shown in brackets, followed by their genus-level assignment. Only CAZyme hits to Glycoside Hydrolases (“GH”) or Polysaccharide Lyases (“PL”) are included in the summarized CAZyme Counts per MAG, and are color coded by their predicted substrate. Broad substrate predictions are based off Berlmont *et al*. 2015. Emboldened MAGs indicate taxa that were active during the demise phase, as indicated by metaproteomics (see Supplementary Figure 8). B) Specific CAZyme HMM hits and dbCAN substrate annotations for *Kordia* (mARW1_16). Hits with no substrate predicted (n=81) were removed for visual purposes.

### Sulfitobacter, an ecologically relevant Roseobacter-clade member, required Kordia to process P. tricornutum lysate

Because *Rhodobacteraceae* taxa present in the demise microbiome generally lacked the capability to degrade complex carbon sources, we hypothesized that the activity of *Kordia* sp, a specialized biopolymer degrader, might allow for the growth of these small molecule users during diatom demise. To test this, we performed a laboratory experiment using a representative *Kordia* strain (*K. algicida* OT1) and an isolate of the *Rhodobacteraceae* family (*Sulfitobacter* sp. N5S). This *Rhodobacteraceae* strain was selected because it is related to relatively abundant MAGs [*Sulfitobacter* sp. (mARW1_117) and *Phaeobacter* sp. (mARW1_12), Figure 3] in the *P. tricornutum* ponds and could be classified as a strict small molecule user due to a lack of CAZymes in its genome (Figure 6A). Further, this genus falls within the Marine Roseobacter Clade, an ecologically relevant and ubiquitous group found in marine environments (61, 62). This isolate also grows in co-culture with *P. tricornutum* for several generations with no added organic matter (Supplementary Figure 10).

**Figure 6.**
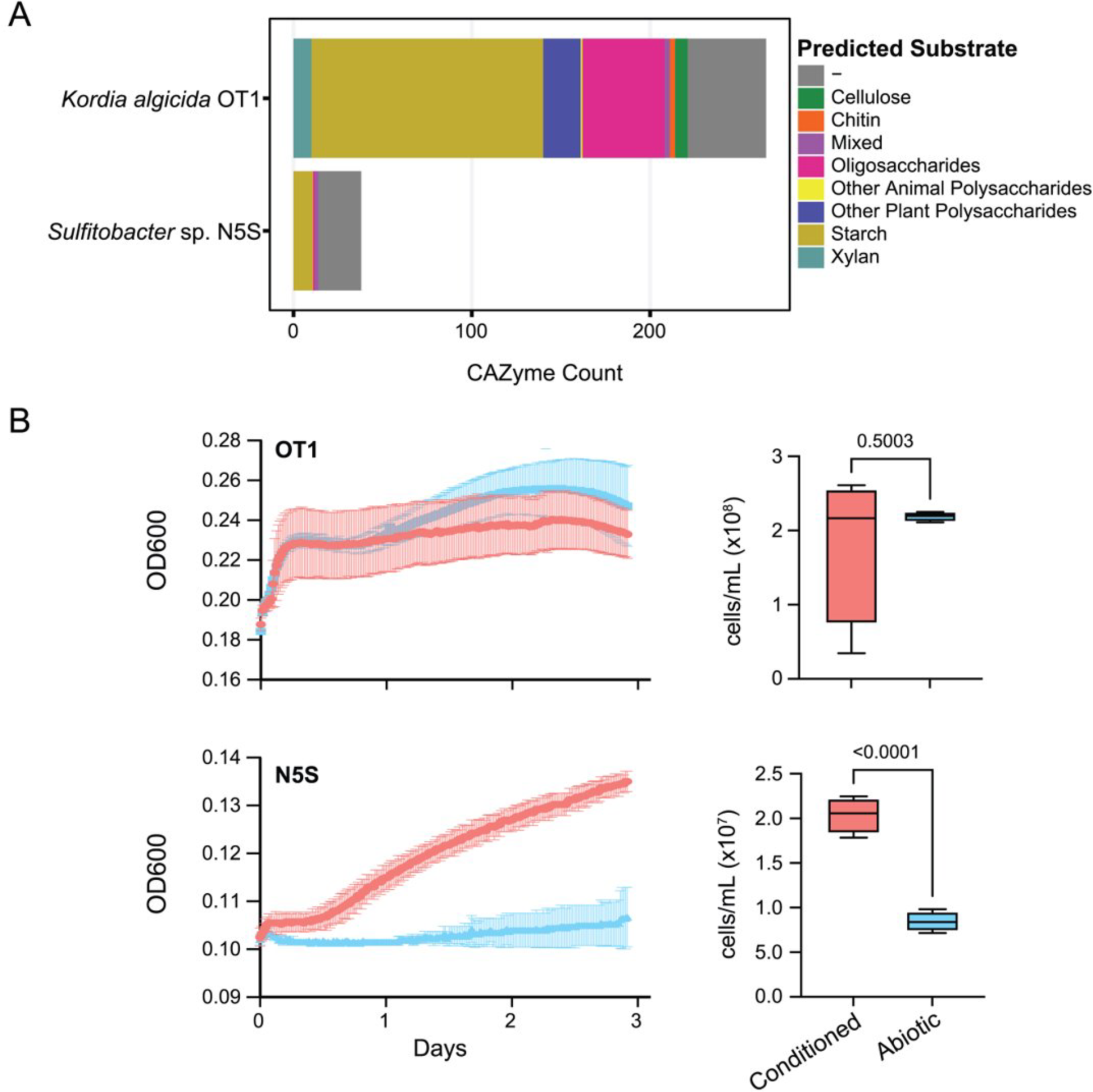
*Sulfitobacter* sp. N5S (a representative *Rhodobacteraceae* strain) requires *Kordia* to pre-condition *P. tricornutum* lysate for growth in culture. A) CAZyme summary of representative lab strains of *Kordia* (*K. algicida* OT1) and a *Rhodobacteraceae* (*Sulfitobacter sp.* N5S) used for the experiment. B) Growth of OT1 *versus* N5S on *P. tricornutum* lysate either pre-conditioned by OT-1 (“Conditioned”) or without any added bacteria (“Abiotic” control). Growth curves show the mean and standard deviation (error bars, n=5) of OD600 values taken every 20 minutes. The boxplots to the right show the cell abundances at the final timepoint (n=4). P-values are generated using unpaired two-tailed *t*-tests.

*K. algicida* OT1 has been shown to have algicidal activity towards *P. tricornutum* (28) and harbors a diverse repertoire of CAZymes targeting complex carbon sources (Figure 6A). So, we grew OT1 on *P. tricornutum* lysate, an analog to the demise phase of the algal ponds for 5 days, and then took that spent media (“Conditioned” lysate) and inoculated it with N5S. We found that N5S was unable to grow in abiotic lysate (lysate incubated in parallel without added cells), but grew in conditioned lysate, with significantly different final cell abundances between the treatments (p<0.0001, Figure 6B). OT1 grew similarly in both its own conditioned lysate and abiotic lysate, with no significant difference in the final cell abundances (p=0.5003, Figure 6B). This suggests N5S was dependent on *Kordia* to access the organic matter present in *P. tricornutum* lysate. Furthermore, the finding that OT1 did not grow differentially under conditioned *versus* abiotic lysate suggests that not all of the algal-derived carbon was fully metabolized during the first incubation. However, our results show this timeframe of conditioning was sufficient to enhance N5S growth.

### Genomic capability to metabolize alternative compounds in marine Rhodobacteraceae

Members of the *Rhodobacteraceae* family are globally abundant in the oceans and well known for having the ability to rapidly respond to increased substrate availability and acquire carbon and energy from alternative sources (61, 63, 64). We hypothesized that the *Rhodobacteraceae* population emerged during the algal demise due to increased labile organic matter released from *Kordia* activity, while persisting under high competition in the growth phase on alternative carbon and energy sources. Thus, we screened all MAGs from the *P. tricornutum* ponds for pathways that confer the ability to utilize aromatic carbohydrates, fix inorganic carbon, and gain energy from light (Figure 7). We found that the ability to degrade aromatic compounds such as 4-hydroxybenzoic acid (4HBA), coumaric acid and phenylacetic acid was common, as was aerobic anoxygenic phototrophy for light-driven metabolism (Figure 7A). However, most notable was that 90-100% of all MAGs in the *Rhodobacteraceae*, *Rhizobiaceae* and *Stappiaceae* families contained the complete set of genes for carbon monoxide dehydrogenase (COX, *coxL* + *coxM* + *coxS*). Moreover, we detected protein expression in the metaproteome of multiple COX subunits across these families, with mARW1_20 (*Nioella sedimensis*), mARW1_25 (unclassified *Rhodobacteraceae*), and mARW7_5 (*Hoeflea* sp.) recruiting all three COX subunits (Figure 7B). The expression patterns of individual COX proteins across timepoints were variable, some having highest relative expression on day 1 (Supplementary Figure 11). Contrary, we detected sparse protein expression of aromatic compound degradation and light-driven metabolism (Supplementary Figure 12). This suggests that carbon monoxide utilization was widespread and active in this group, especially during the growth phase.

**Figure 7.**
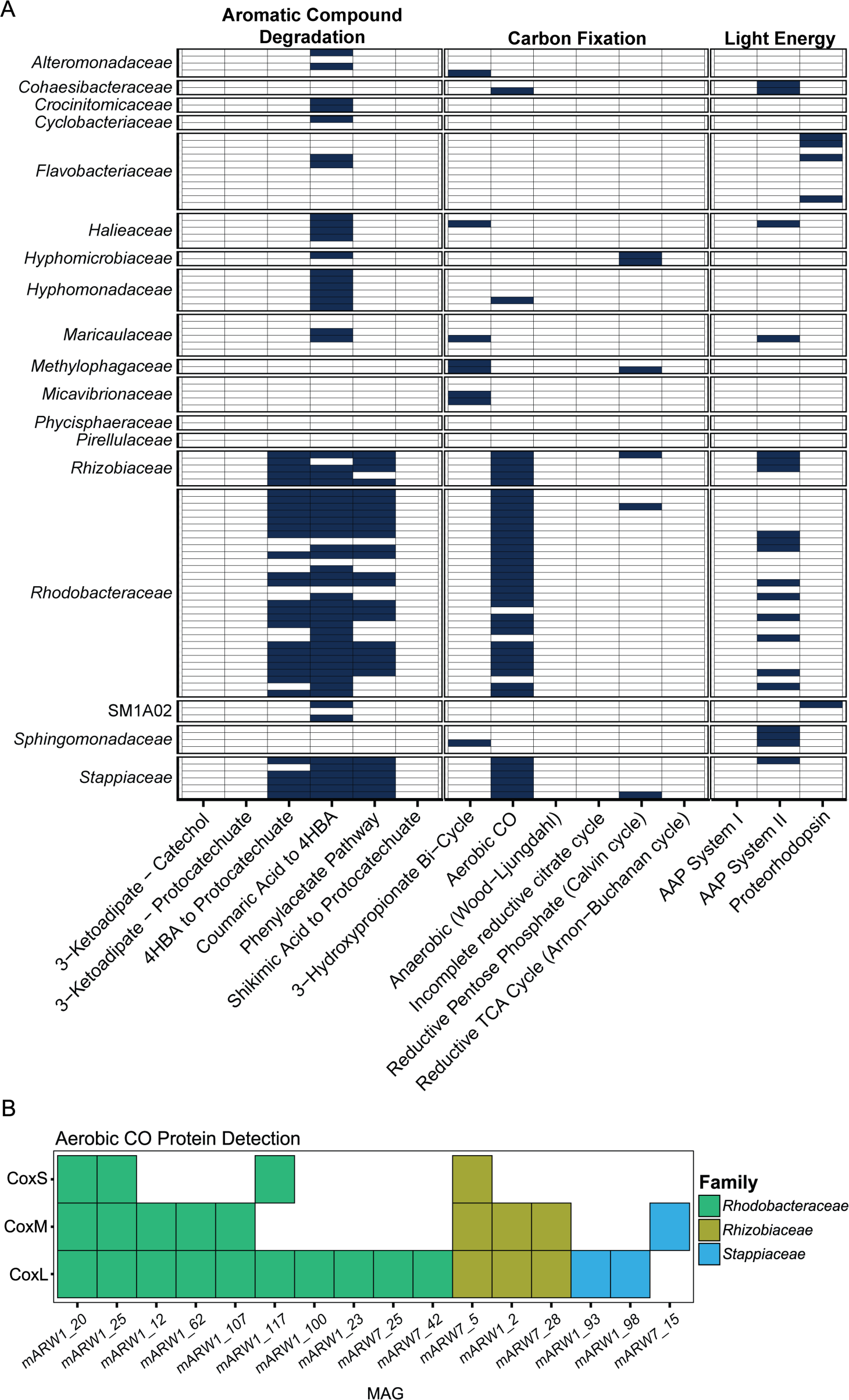
Carbon monoxide (CO) oxidation as a potential alternative carbon and energy generating mechanism used by *Rhodobacteraceae* in algal ponds. A) Metabolic potential across all P. tricornutum pond MAGs to use alternative carbon and energy sources. All screened pathways are shown (x-axis), where dark blue tiles indicate 100% of pathway detection (i.e., all known genes involved in that pathway were detected) per-MAG (y-axis, grouped by family). B) Protein presence/absence of each CODH subunit across MAGs in the metaproteome. Only MAGs that had a detected protein mapped to CODH out of 13 samples are shown.

## Discussion

Our study documents a serendipitous algal population crash that occurred during a multi-omics assessment of outdoor *P. tricornutum* ponds. Initially, we set out to characterize the microbiome dynamics of two algal species cultivated in biofuel ponds. Despite close proximity, distinct microbiomes were observed, shown to be driven by host-specific physiology (65). However, a drop in bacterial diversity during *P. tricornutum* demise corresponded with a “bloom” of *Kordia*, a genus known to be algicidal against *P. tricornutum* (28). We discuss 1) features specific to *Kordia* that potentially led to its takeover 2) how *Kordia* plays an indirect role in shaping the microbiome, and 3) unique metabolic features of downstream microbiota that may explain their emergence during algal demise. We propose cross-feeding as a key determinant shaping the microbiome of algae, specifically between biopolymer degraders like *Kordia* (*Flavobacteriaceae*) and copiotrophic *Rhodobacteraceae* during algal demise.

The *Kordia*-specific proteomics analysis collectively suggests antagonistic interactions with other bacteria and/or *P. tricornutum*. Currently, the literature indicates that *Kordia* secretes a protease involved in algal killing (28). Our proteomics dataset only covered intracellular proteins, limiting robust detection of extracellular proteases involved in algal lysis. However, we found that protease/peptidase-like proteins comprised a major fraction of the *Kordia* proteome during demise. These proteins have widely ranging substrate specificity and roles [*e.g.,* protein folding, biopolymer breakdown, pathogenesis; (66)] and particular algicidal proteases could be species-specific which complicates comparisons with previous studies to validate algicidal activity (67).

Two of the most abundant *Kordia* proteins belonged to pathways putatively involved in algal- or bacterial antagonism. The first was a putative lysine oxidase (LodA, aka marinocine), a protein shown to be antimicrobial *via* hydrogen peroxide (H_2_O_2_) produced during amino acid deamination (68). LodA-driven H_2_O_2_ production can lead to bactericidal behavior between species under competition (68–70). Heretofore, an L-amino acid oxidase produced by a bacterium has only been shown to be algicidal against the cyanobacterium *Microcystis aeruginosa* (71). Direct production of H_2_O_2_ *via* LodA by a bacterium against eukaryotes has not been shown. However, asparagine addition to *P. tricornutum* cultures triggered cellular death due to H_2_O_2_ production (72). Analysis of the *P. tricornutum* genome identified an ortholog to the confirmed H_2_O_2_-producing L-amino acid oxidase (LAAO) of *Chlamydomonas reinhardtii* and other algae (72). Second, several components of a type VI secretion system (T6SS) gene cluster were expressed, one being among the fourth most abundant *Kordia* proteins. T6SS is involved in anti-bacterial and eukaryotic defense (73–75), a potential strategy to reduce competition or gain limiting resources (76). It’s possible that *Kordia* employed T6SS activity against *P. tricornutum*, however T6SS-guided algal killing has not yet been investigated (76).

Despite the absence of causal evidence linking *Kordia* to *P. tricornutum* demise, *Kordia’s* bloom during algal demise may be attributed to its ability to access diatom-derived biopolymers. *Kordia* is a *Flavobacteriaceae* (phylum Bacteroidota), a group known to degrade and consume biopolymers (77, 78). Similar to a previous *Kordia* genomic analysis (79), we found many polysaccharide-degrading CAZymes (80) in the *Kordia* MAG. Furthermore, the most abundant *Kordia* proteins detected were assigned to several polysaccharide utilization loci systems used to extracellularly hydrolyze and import biopolymer carbon (24, 78). Although the substrates cannot currently be determined based on sequences alone, these proteins sit upstream of expressed CAZymes putatively targeting beta-glucans commonly found in diatom cell walls (81). Overall, these observations supported our initial hypothesis that biopolymer degradation and the *Flavobacteriaceae* would be prevalent during algal decay (14, 17, 18).

During algal demise, the diversity of other putative biopolymer degraders present initially during growth was reduced, and only one other *Flavobacteriaceae* (*Muricauda* sp.) remained. We hypothesize that *Kordia* sp. removed competing bacteria, either by being optimized to demise conditions, and/or by antibacterial modes (*e.g.,* LodA and T6SS). However, several distinct genera of the *Rhodobacteraceae* co-occurred with *Kordia*, some assigned to the Marine Roseobacter Clade [MRC; *e.g., Roseovarius* and *Phaeobacter*; (61, 62)]. The emergence of distinct *Rhodobacteraceae* during demise (compared to growth) suggests these specific genera may be suited for this niche space. Significant genomic variation within the *Rhodobacteraceae* may confer adaptation to the widely ranging conditions along algal blooms (14, 18, 19). Nevertheless, the *Rhodobacteraceae* commonly specialize in the uptake of algal-DOC released during growth and development (20, 22). Specifically, *Rhodobacteraceae* taxa are enriched in transporters for dicarboxylic acids, sugars, dimethylsulfoniopropionate, and polyamines (63, 82, 83)]. Thus, our analysis of polysaccharide CAZymes did not indicate any unique divergence from typical *Rhodobacteraceae* (63, 84, 85) to access carbon directly from biopolymers enriched during algal demise. The appearance of distinct *Rhodobacteraceae* co-occurring with *Kordia* was intriguing, as we initially hypothesized they would be more diverse during algal growth.

Instead, our results collectively indicate that the co-occurrence of distinct *Rhodobacteraceae* with *Kordia* sp. during algal demise may be a consequence of cross-feeding. Cross-feeding is a ubiquitous phenomenon that structures microbial communities (86), and previous studies have speculated that cross-feeding between *Rhodobacteraceae* and *Flavobacteriaceae* led to transparent exopolymer particle degradation within a diatom bloom (87). When biopolymers are abundant, community assembly generally follows primary “degraders” that initially colonize these resources (88, 89). Extracellular biopolymer hydrolysis releases oligo- and monomers forms into the environment, which are consumed by “exploiters” (direct uptake) and/or “scavengers” [uptake of exuded metabolites; (88, 90–92)]. Our laboratory experiment supports this idea by demonstrating that the pre-processing of *P. tricornutum* lysate by *Kordia* (degrader) allowed the growth of a representative MRC strain (*Sulfitobacter* sp. N5S) under conditions analogous to algal demise. It’s not known whether N5S benefitted from direct breakdown products, other exuded or released metabolites, or antimicrobial compound reduction. Yet, these results demonstrate how interspecies (inter-family in this case) interactions, driven by *Kordia*, can structure the microbiome. Previously, addition of *Kordia algicida* to a natural phytoplankton community removed dominant species, released algal nutrients *via* lysis, and possibly enhanced DOC *via* biopolymer degradation (93). It’s possible that *Rhodobacteraceae* populations were exploiting diatom-stored nutrients released *via* lysis in the ARW ponds, however our experimental system suggests N5S required *Kordia* conditioning to access DOC. A follow up experiment might involve testing this hypothesis under mixed culture conditions (92) to investigate ecological interactions between *Kordia* and *Rhodobacteraceae*. Further, the structuring of microbiota across the algal life cycle in general likely reflects different cross-feeding networks between microbes that both produce and consume one another’s metabolites and is not necessarily restricted to algal demise conditions.

Lastly, we probed the metabolic potential of the *Rhodobacteraceae* populations in the ponds. Although many *Rhodobacteraceae* taxa emerged during algal demise, other *Rhodobacteraceae* were also present during the *P. tricornutum* growth stage, alluding to this group’s “generalist” lifestyle. Oxidation of the trace gas, carbon monoxide (CO), can generate supplementary energy or carbon for persistence under “survival” conditions when other sources are limited (94, 95). For MRC members, the CO oxidation pathway is common and hypothesized to be used primarily for energy due to a lack of carbon fixation pathways (63). Here, we propose that CO may have been a viable source of energy, and potentially carbon, for the *Rhodobacteraceae* populations in the ponds. Indeed, a previous proteomic assessment done by our research group on *Rhodophyticola* sp. PT6CLA (*Rhodobacteraceae*) co-cultured with *P. tricornutum* in the laboratory revealed that CoxMSL was a highly expressed proteins cluster (11), suggesting this may be a common biogeochemical transformation occurring in algal microbiomes. Further, CO oxidation may contribute significantly to carbon flow in algal ponds yet has not been investigated in algal pond microbiomes. Additionally, aromatic compound degradation pathways were also widespread among the *Rhodobacteraceae* MAGs, specifically for the degradation of coumaric acid to 4-hydroxybenzoate (4HBA) and 4HBA to protocatechuate, aromatics previously found to be exuded in abundance by *P. tricornutum* (16). Overall, these observations collectively emphasize the role of flexible carbon and energy metabolisms in structuring the microbiome within algal ponds and provide insight into potential survival mechanisms used by the *Rhodobacteraceae* when under competition for resources.

## Conclusions

Our characterization of a diatom population crash in outdoor ponds suggests *Kordia* sp. is both an environmentally and industrially relevant organism that can structure algal and bacterial communities. Overall, our analysis provides insight into potential virulence pathways employed by *Kordia* that could have led to *P. tricornutum* demise and/or removal of bacterial competitors. We showed how cross-feeding interactions could be an indirect consequence of *Kordia* infection in the algal microbiome. This is relevant to biofuel pond management, where the application of a primary degrader is followed with small-molecule remineralizers to remove remaining organic matter following pond crashes. Lastly, we highlight features of the globally relevant *Rhodobacteraceae* population, suggesting they are well adapted to respond to rapid changes in algal growth and sustain abundances using alternative carbon and energy sources.

## Acknowledgements

We thank D. Nilson, S. Mabery, M. Hwang, C. Swink, and J. Wollard for laboratory assistance. We also thank A. Navid and J. Casey for their valuable discussions on the work. This work was supported by the US Department of Energy’s (DOE) Genomic Science Program through the LLNL mBioSpheres Science Focus Area grant # SCW1039 and carried out at Lawrence Livermore National Laboratory (LLNL) under Contract DE-AC52-07NA27344. Metagenomics and ribosomal RNA amplicon sequencing was carried out at the Joint Genome Institute (JGI) through Community Sequencing Program award # 1939 (received by X. M.). JGI is sponsored by the DOE’s Biological and Environmental Research program and operated under Contract Nos. DE-AC02-05CH11231. Proteomic data was collected as part of project award 50220 (received by T.J.S.) under the FICUS program and used resources at the Environmental Molecular Sciences Laboratory (EMSL), which is a DOE Office of Science User Facilities and operated under contract DE-AC05-76RL01830. Institution Paper Number LLNL-JRNL-865370-DRAFT

## References

1. Cole JJ. Interactions between bacteria and algae in aquatic ecosystems. Annual review of Ecology and systematics. 1982;13(1):291–314.

2. Seymour JR, Amin SA, Raina J-B, Stocker R. Zooming in on the phycosphere: the ecological interface for phytoplankton–bacteria relationships. Nature microbiology. 2017;2(7):1–12.

3. Cirri E, Pohnert G. Algae− bacteria interactions that balance the planktonic microbiome. New Phytologist. 2019;223(1):100–6.

4. Kuhlisch C, Shemi A, Barak-Gavish N, Schatz D, Vardi A. Algal blooms in the ocean: hot spots for chemically mediated microbial interactions. Nature Reviews Microbiology. 2023:1–17.

5. Wheeler PA, Kirchman DL. Utilization of inorganic and organic nitrogen by bacteria in marine systems 1. Limnology and Oceanography. 1986;31(5):998–1009.

6. Croft MT, Lawrence AD, Raux-Deery E, Warren MJ, Smith AG. Algae acquire vitamin B12 through a symbiotic relationship with bacteria. Nature. 2005;438(7064):90–3.

7. De-Bashan LE, Antoun H, Bashan Y. Involvement of indole-3-acetic acid produced by the growth-promoting bacterium Azospirillum spp. in promoting growth of chlorella vulgaris 1. Journal of Phycology. 2008;44(4):938–47.

8. Amin SA, Green DH, Hart MC, Küpper FC, Sunda WG, Carrano CJ. Photolysis of iron– siderophore chelates promotes bacterial–algal mutualism. Proceedings of the National Academy of Sciences. 2009;106(40):17071–6.

9. Amin S, Hmelo L, Van Tol H, Durham B, Carlson L, Heal K, et al. Interaction and signalling between a cosmopolitan phytoplankton and associated bacteria. Nature. 2015;522(7554):98–101.

10. Ramanan R, Kim B-H, Cho D-H, Oh H-M, Kim H-S. Algae–bacteria interactions: evolution, ecology and emerging applications. Biotechnology advances. 2016;34(1):14–29.

11. Mayali X, Samo TJ, Kimbrel JA, Morris MM, Rolison K, Swink C, et al. Single-cell isotope tracing reveals functional guilds of bacteria associated with the diatom Phaeodactylum tricornutum. Nature Communications. 2023;14(1):5642.

12. Mayali X, Azam F. Algicidal bacteria in the sea and their impact on algal blooms 1. Journal of Eukaryotic Microbiology. 2004;51(2):139–44.

13. Meyer N, Bigalke A, Kaulfuß A, Pohnert G. Strategies and ecological roles of algicidal bacteria. FEMS Microbiology Reviews. 2017;41(6):880–99.

14. Buchan A, LeCleir GR, Gulvik CA, González JM. Master recyclers: features and functions of bacteria associated with phytoplankton blooms. Nature Reviews Microbiology. 2014;12(10):686–98.

15. Ferrer-González FX, Widner B, Holderman NR, Glushka J, Edison AS, Kujawinski EB, et al. Resource partitioning of phytoplankton metabolites that support bacterial heterotrophy. The ISME journal. 2021;15(3):762–73.

16. Brisson V, Swink C, Kimbrel J, Mayali X, Samo T, Kosina SM, et al. Dynamic Phaeodactylum tricornutum exometabolites shape surrounding bacterial communities. New Phytologist. 2023;239(4):1420–33.

17. Riemann L, Steward GF, Azam F. Dynamics of bacterial community composition and activity during a mesocosm diatom bloom. Applied and environmental microbiology. 2000;66(2):578–87.

18. Pinhassi J, Sala MM, Havskum H, Peters F, Guadayol O, Malits A, et al. Changes in bacterioplankton composition under different phytoplankton regimens. Applied and Environmental Microbiology. 2004;70(11):6753–66.

19. Teeling H, Fuchs BM, Becher D, Klockow C, Gardebrecht A, Bennke CM, et al. Substrate-controlled succession of marine bacterioplankton populations induced by a phytoplankton bloom. Science. 2012;336(6081):608–11.

20. Pontiller B, Martínez-García S, Joglar V, Amnebrink D, Pérez-Martínez C, González JM, et al. Rapid bacterioplankton transcription cascades regulate organic matter utilization during phytoplankton bloom progression in a coastal upwelling system. The ISME Journal. 2022;16(10):2360–72.

21. Hellebust JA. Excretion of some organic compounds by marine phytoplankton 1. Limnology and Oceanography. 1965;10(2):192–206.

22. Kieft B, Li Z, Bryson S, Hettich RL, Pan C, Mayali X, et al. Phytoplankton exudates and lysates support distinct microbial consortia with specialized metabolic and ecophysiological traits. Proceedings of the national academy of sciences. 2021;118(41):e2101178118.

23. Biddanda B, Benner R. Carbon, nitrogen, and carbohydrate fluxes during the production of particulate and dissolved organic matter by marine phytoplankton. Limnology and Oceanography. 1997;42(3):506–18.

24. McKee LS, La Rosa SL, Westereng B, Eijsink VG, Pope PB, Larsbrink J. Polysaccharide degradation by the Bacteroidetes: mechanisms and nomenclature. Environmental Microbiology Reports. 2021;13(5):559–81.

25. Chisti Y. Large-scale production of algal biomass: raceway ponds. Algae biotechnology: Products and processes. 2016:21–40.

26. Lane TW. Barriers to microalgal mass cultivation. Current Opinion in Biotechnology. 2022;73:323–8.

27. Carney LT, Lane TW. Parasites in algae mass culture. Frontiers in microbiology. 2014;5:278.

28. Paul C, Pohnert G. Interactions of the algicidal bacterium Kordia algicida with diatoms: regulated protease excretion for specific algal lysis. PloS one. 2011;6(6):e21032.

29. Parada AE, Needham DM, Fuhrman JA. Every base matters: assessing small subunit rRNA primers for marine microbiomes with mock communities, time series and global field samples. Environmental microbiology. 2016;18(5):1403–14.

30. Apprill A, McNally S, Parsons R, Weber L. Minor revision to V4 region SSU rRNA 806R gene primer greatly increases detection of SAR11 bacterioplankton. Aquatic Microbial Ecology. 2015;75(2):129–37.

31. Stoeck T, Bass D, Nebel M, Christen R, Jones MD, Breiner HW, et al. Multiple marker parallel tag environmental DNA sequencing reveals a highly complex eukaryotic community in marine anoxic water. Molecular ecology. 2010;19:21–31.

32. Callahan BJ, McMurdie PJ, Rosen MJ, Han AW, Johnson AJA, Holmes SP. DADA2: High-resolution sample inference from Illumina amplicon data. Nature methods. 2016;13(7):581–3.

33. Quast C, Pruesse E, Yilmaz P, Gerken J, Schweer T, Yarza P, et al. The SILVA ribosomal RNA gene database project: improved data processing and web-based tools. Nucleic acids research. 2012;41(D1):D590–D6.

34. Wang Q, Garrity GM, Tiedje JM, Cole JR. Naive Bayesian classifier for rapid assignment of rRNA sequences into the new bacterial taxonomy. Applied and environmental microbiology. 2007;73(16):5261–7.

35. R Core Team R. R: A language and environment for statistical computing. 2013.

36. Wilkinson L. ggplot2: elegant graphics for data analysis by Wickham, H. Oxford University Press; 2011.

37. Lin H, Peddada S. Analysis of compositions of microbiomes with bias correction. Nat Commun 11: 3514. 2020.

38. Dixon P. VEGAN, a package of R functions for community ecology. Journal of vegetation science. 2003;14(6):927–30.

39. Huntemann M, Ivanova NN, Mavromatis K, Tripp HJ, Paez-Espino D, Tennessen K, et al. The standard operating procedure of the DOE-JGI Metagenome Annotation Pipeline (MAP v.4). Standards in Genomic Sciences. 2016;11(1):17.

40. Nurk S, Meleshko D, Korobeynikov A, Pevzner PA. metaSPAdes: a new versatile metagenomic assembler. Genome research. 2017;27(5):824–34.

41. Uritskiy GV, DiRuggiero J, Taylor J. MetaWRAP—a flexible pipeline for genome-resolved metagenomic data analysis. Microbiome. 2018;6:1–13.

42. Parks DH, Imelfort M, Skennerton CT, Hugenholtz P, Tyson GW. CheckM: assessing the quality of microbial genomes recovered from isolates, single cells, and metagenomes. Genome research. 2015;25(7):1043–55.

43. Prjibelski A, Antipov D, Meleshko D, Lapidus A, Korobeynikov A. Using SPAdes de novo assembler. Current protocols in bioinformatics. 2020;70(1):e102.

44. Olm MR, Brown CT, Brooks B, Banfield JF. dRep: a tool for fast and accurate genomic comparisons that enables improved genome recovery from metagenomes through de-replication. The ISME journal. 2017;11(12):2864–8.

45. Bushnell B. BBTools software packag. e. 2014.

46. Morris MM, Kimbrel JA, Geng H, Tran-Gyamfi MB, Yu ET, Sale KL, et al. Bacterial Community Assembly, Succession, and Metabolic Function during Outdoor Cultivation of *Microchloropsis salina*. mSphere. 2022;7(4):e00231–22.

47. Chaumeil P-A, Mussig AJ, Hugenholtz P, Parks DH. GTDB-Tk: a toolkit to classify genomes with the Genome Taxonomy Database. Oxford University Press; 2020.

48. Brettin T, Davis JJ, Disz T, Edwards RA, Gerdes S, Olsen GJ, et al. RASTtk: a modular and extensible implementation of the RAST algorithm for building custom annotation pipelines and annotating batches of genomes. Scientific reports. 2015;5(1):1–6.

49. Snyder E, Kampanya N, Lu J, Nordberg EK, Karur H, Shukla M, et al. PATRIC: the VBI pathosystems resource integration center. Nucleic acids research. 2007;35(suppl_1):D401–D6.

50. Zhang H, Yohe T, Huang L, Entwistle S, Wu P, Yang Z, et al. dbCAN2: a meta server for automated carbohydrate-active enzyme annotation. Nucleic acids research. 2018;46(W1):W95–W101.

51. Emms DM, Kelly S. OrthoFinder: phylogenetic orthology inference for comparative genomics. Genome biology. 2019;20:1–14.

52. Emms DM, Kelly S. OrthoFinder: solving fundamental biases in whole genome comparisons dramatically improves orthogroup inference accuracy. Genome biology. 2015;16:1–14.

53. Emms DM, Kelly S. STRIDE: species tree root inference from gene duplication events. Molecular biology and evolution. 2017;34(12):3267–78.

54. Letunic I, Bork P. Interactive Tree Of Life (iTOL) v5: an online tool for phylogenetic tree display and annotation. Nucleic acids research. 2021;49(W1):W293–W6.

55. Bowler C, Allen AE, Badger JH, Grimwood J, Jabbari K, Kuo A, et al. The Phaeodactylum genome reveals the evolutionary history of diatom genomes. Nature. 2008;456(7219):239–44.

56. Kim S, Pevzner PA. MS-GF+ makes progress towards a universal database search tool for proteomics. Nature communications. 2014;5(1):5277.

57. Monroe ME, Shaw JL, Daly DS, Adkins JN, Smith RD. MASIC: a software program for fast quantitation and flexible visualization of chromatographic profiles from detected LC–MS (/MS) features. Computational biology and chemistry. 2008;32(3):215–7.

58. Florens L, Carozza MJ, Swanson SK, Fournier M, Coleman MK, Workman JL, et al. Analyzing chromatin remodeling complexes using shotgun proteomics and normalized spectral abundance factors. Methods. 2006;40(4):303–11.

59. Samo TJ, Kimbrel JA, Nilson DJ, Pett-Ridge J, Weber PK, Mayali X. Attachment between heterotrophic bacteria and microalgae influences symbiotic microscale interactions. Environmental microbiology. 2018;20(12):4385–400.

60. Lucas-Elío P, Gómez D, Solano F, Sanchez-Amat A. The antimicrobial activity of marinocine, synthesized by Marinomonas mediterranea, is due to hydrogen peroxide generated by its lysine oxidase activity. Journal of bacteriology. 2006;188(7):2493–501.

61. Buchan A, González JM, Moran MA. Overview of the marine Roseobacter lineage. Applied and environmental microbiology. 2005;71(10):5665–77.

62. Luo H, Moran MA. Evolutionary ecology of the marine Roseobacter clade. Microbiology and Molecular Biology Reviews. 2014;78(4):573–87.

63. Moran MA, Belas R, Schell M, González J, Sun F, Sun S, et al. Ecological genomics of marine Roseobacters. Applied and environmental microbiology. 2007;73(14):4559–69.

64. Wagner-Döbler I, Biebl H. Environmental biology of the marine Roseobacter lineage. Annu Rev Microbiol. 2006;60:255–80.

65. Kimbrel JA, Samo TJ, Ward C, Nilson D, Thelen MP, Siccardi A, et al. Host selection and stochastic effects influence bacterial community assembly on the microalgal phycosphere. Algal Research. 2019;40:101489.

66. Rao MB, Tanksale AM, Ghatge MS, Deshpande VV. Molecular and biotechnological aspects of microbial proteases. Microbiology and molecular biology reviews. 1998;62(3):597–635.

67. Syhapanha KS, Russo DA, Deng Y, Meyer N, Poulin RX, Pohnert G. Transcriptomics-guided identification of an algicidal protease of the marine bacterium Kordia algicida OT-1. MicrobiologyOpen. 2023;12(5):e1387.

68. Lucas-Elio P, Hernandez P, Sanchez-Amat A, Solano F. Purification and partial characterization of marinocine, a new broad-spectrum antibacterial protein produced by Marinomonas mediterranea. Biochimica et Biophysica Acta (BBA)-General Subjects. 2005;1721(1-3):193–203.

69. Mai-Prochnow A, Lucas-Elio P, Egan S, Thomas T, Webb JS, Sanchez-Amat A, et al. Hydrogen peroxide linked to lysine oxidase activity facilitates biofilm differentiation and dispersal in several gram-negative bacteria. Journal of bacteriology. 2008;190(15):5493–501.

70. Tong H, Chen W, Shi W, Qi F, Dong X. SO-LAAO, a novel L-amino acid oxidase that enables Streptococcus oligofermentans to outcompete Streptococcus mutans by generating H2O2 from peptone. Journal of Bacteriology. 2008;190(13):4716–21.

71. Chen WM, Sheu FS, Sheu SY. Novel L-amino acid oxidase with algicidal activity against toxic cyanobacterium Microcystis aeruginosa synthesized by a bacterium Aquimarina sp. Enzyme and microbial technology. 2011;49(4):372–9.

72. Contreras JA, Gillard JT. Asparagine-based production of hydrogen peroxide triggers cell death in the diatom Phaeodactylum tricornutum. Botany Letters. 2021;168(1):6–17.

73. Ray A, Schwartz N, de Souza Santos M, Zhang J, Orth K, Salomon D. Type VI secretion system MIX-effectors carry both antibacterial and anti-eukaryotic activities. EMBO reports. 2017;18(11):1978–90.

74. Jiang F, Waterfield NR, Yang J, Yang G, Jin Q. A Pseudomonas aeruginosa type VI secretion phospholipase D effector targets both prokaryotic and eukaryotic cells. Cell host & microbe. 2014;15(5):600–10.

75. Coulthurst S. The Type VI secretion system: a versatile bacterial weapon. Microbiology. 2019;165(5):503–15.

76. Kempnich MW, Sison-Mangus MP. Presence and abundance of bacteria with the Type VI secretion system in a coastal environment and in the global oceans. Plos one. 2020;15(12):e0244217.

77. Fernández-Gómez B, Richter M, Schüler M, Pinhassi J, Acinas SG, González JM, et al. Ecology of marine Bacteroidetes: a comparative genomics approach. The ISME journal. 2013;7(5):1026–37.

78. Grondin JM, Tamura K, Déjean G, Abbott DW, Brumer H. Polysaccharide utilization loci: fueling microbial communities. Journal of bacteriology. 2017;199(15):10.1128/jb.00860-16.

79. Lim Y, Kang I, Cho J-C. Genome characteristics of Kordia antarctica IMCC3317T and comparative genome analysis of the genus Kordia. Scientific Reports. 2020;10(1):14715.

80. Huang L, Zhang H, Wu P, Entwistle S, Li X, Yohe T, et al. dbCAN-seq: a database of carbohydrate-active enzyme (CAZyme) sequence and annotation. Nucleic Acids Research. 2018;46(D1):D516–D21.

81. Myklestad SM. Production, chemical structure, metabolism, and biological function of the (1→ 3)-linked, β3-D-glucans in diatoms. Biological oceanography. 1989;6(3-4):313–26.

82. Newton RJ, Griffin LE, Bowles KM, Meile C, Gifford S, Givens CE, et al. Genome characteristics of a generalist marine bacterial lineage. The ISME journal. 2010;4(6):784–98.

83. Poretsky RS, Sun S, Mou X, Moran MA. Transporter genes expressed by coastal bacterioplankton in response to dissolved organic carbon. Environmental microbiology. 2010;12(3):616–27.

84. Cha Q-Q, Liu S-S, Dang Y-R, Ren X-B, Xu F, Li P-Y, et al. Ecological function and interaction of different bacterial groups during alginate processing in coastal seawater community. Environment International. 2023;182:108325.

85. Priest T, Vidal-Melgosa S, Hehemann J-H, Amann R, Fuchs BM. Carbohydrates and carbohydrate degradation gene abundance and transcription in Atlantic waters of the Arctic. ISME communications. 2023;3(1):130.

86. D’Souza G, Shitut S, Preussger D, Yousif G, Waschina S, Kost C. Ecology and evolution of metabolic cross-feeding interactions in bacteria. Natural Product Reports. 2018;35(5):455–88.

87. Taylor JD, Cottingham SD, Billinge J, Cunliffe M. Seasonal microbial community dynamics correlate with phytoplankton-derived polysaccharides in surface coastal waters. The ISME journal. 2014;8(1):245–8.

88. Enke TN, Datta MS, Schwartzman J, Cermak N, Schmitz D, Barrere J, et al. Modular assembly of polysaccharide-degrading marine microbial communities. Current Biology. 2019;29(9):1528–35. e6.

89. Pontrelli S, Szabo R, Pollak S, Schwartzman J, Ledezma-Tejeida D, Cordero OX, et al. Metabolic cross-feeding structures the assembly of polysaccharide degrading communities. Science advances. 2022;8(8):eabk3076.

90. Cordero OX, Datta MS. Microbial interactions and community assembly at microscales. Current opinion in microbiology. 2016;31:227–34.

91. Datta MS, Sliwerska E, Gore J, Polz MF, Cordero OX. Microbial interactions lead to rapid micro-scale successions on model marine particles. Nature communications. 2016;7(1):11965.

92. D’souza G, Schwartzman J, Keegstra J, Schreier JE, Daniels M, Cordero O, et al. Interspecies interactions determine growth dynamics of biopolymer degrading populations in microbial communities. bioRxiv. 2023:2023.03. 22.533748.

93. Bigalke A, Meyer N, Papanikolopoulou LA, Wiltshire KH, Pohnert G. The algicidal bacterium Kordia algicida shapes a natural plankton community. Applied and Environmental Microbiology. 2019;85(7):e02779–18.

94. King GM, Weber CF. Distribution, diversity and ecology of aerobic CO-oxidizing bacteria. Nature Reviews Microbiology. 2007;5(2):107–18.

95. Cordero PR, Bayly K, Man Leung P, Huang C, Islam ZF, Schittenhelm RB, et al. Atmospheric carbon monoxide oxidation is a widespread mechanism supporting microbial survival. The ISME journal. 2019;13(11):2868–81.

