## Supplementary Information for "An algicidal bacterium shapes the microbiome during outdoor diatom cultivation collapse"

**Supporting Information**
Article Title: An algicidal bacterium shapes the microbiome during diatom demise

**Authors**
Naomi E. Gilbert Jeffery A. Kimbrel, Ty J. Samo, Anthony J. Siccardi, Rhona K. Stuart[1] Xavier Mayali

### Supplementary Methods

16S/18S analysis

The 16S and 18S rRNA genes were amplified with 16S rRNA V4 primers 515F [GTGYCAGCMGCCGCGGTAA, (33)] and 806R [GGACTACNVGGGTWTCTAAT, (34)], and 18S rRNA V4 primers 565F (CCAGCASCYGCGGTAATTCC) and 908R (ACTTTCGTTCTTGATYRA) (35). Paired reads (2x301 base pair) were processed with DADA2 v1.6.0 (36) for filtering (parameters maxN = 0, maxEE = c(2, 2), truncQ = 2). For the 16S reads, the forward and reverse reads were trimmed to 15-230 and 20-190, respectively. For the 18S data, both the forward and reverse reads were trimmed from 5-275. Chimeric sequences were predicted *de novo* and removed with the removeBimeraDenovo() function in DADA2 using the “consensus” method. 16S and 18S amplicon sequence variants (ASVs) were assigned taxonomy with Silva v132 (37), and the 16S ASVs were further assigned with the RDP classifier v2.11 (38) against training set 16. The taxonomy datasets were filtered to remove ASVs with a “chloroplast” or “mitochondria” family assignment from the 16S dataset.

*Metaproteomics collection and extraction*

SUPOR filter pieces were transferred to 2 mL snap-cap centrifuge tubes (Eppendorf, Hamburg, Germany) with 0.1 mm zirconia beads and bead beat in a Bullet Blender (Next Advance, Averill Park, NY) at speed 8 for 3 minutes at 4°C. After bead beating the lysate was spun into a 15 mL Falcon tube at 2000 xg for 10 min at 4°C. Each sample was transferred to new tubes and a bicinchoninic acid (BCA) assay (Thermo Scientific, Waltham, MA USA) was performed to determine protein concentration. Urea was added to each tube to bring the concentration to 8 M (Sigma-Aldrich, Saint Louis, MO) and dithiothreitol (DTT) was added to the samples at 10 mM. The samples were incubated at 60°C for 30 min with constant shaking at 800 rpm. Samples were then diluted 8-fold for preparation for digestion with 100 mM $NH_4HCO_3$, 1 mM $CaCl_2$ and sequencing-grade modified porcine trypsin (Promega, Madison, WI) was added to all protein samples at a 1:50 (w/w) trypsin-to-protein ratio for 3 h at 37°C. Digested samples were desalted using a 4-probe positive pressure Gilson GX-274 ASPEC™ system (Gilson Inc., Middleton, WI) with Discovery C18 100 mg/1 mL solid phase extraction tubes (Supelco, St. Louis, MO), using the following protocol: 3 mL of methanol was added for conditioning followed by 2 mL of 0.1 % TFA in $H_2O$. The samples were then loaded onto each column followed by 4 mL of 95:5: H2O:I, 0.1 % TFA.

Columns were packed in-house using 360 µm o.d. fused silica (Polymicro Technologies Inc., Phoenix, AZ) with 2-mm sol-gel frits for media retention and contained Jupiter C18 media (Phenomenex, Torrence, CA) in 5 µm particle size for the trapping column (150 µm i.d. x 4 cm long) and 3 µm particle size for the analytical column (75 µm i.d. x 70 cm long). Mobile phases consisted of (A) 0.1 % formic acid in water and (B) 0.1 % formic acid in acetonitrile with the following gradient profile (min, %B): 0, 1; 2, 8; 20, 12; 75, 30; 97, 45; 100, 95; 110, 95; 115, 1; 150, 1. Samples were eluted with 1 mL 80:20 I:H2O, 0.1 % TFA. The samples were concentrated down to ~100 µL using a Speed Vac and a final BCA was performed to determine the peptide concentration and samples were diluted to 0.1 µg/µL with nanopure water for MS analysis.

MS analysis was performed using a Q-Exactive HF mass spectrometer (Thermo Scientific, San Jose, CA) outfitted with a home-made nano-electrospray ionization interface. Electrospray emitters were prepared using 150 um o.d. x 20 um i.d. chemically etched fused silica (58). The ion transfer tube temperature and spray voltage were 320 ºC and 2.2 kV, respectively. Data were collected for 120 min following a 20 min delay from sample injection. FT-MS spectra were acquired from 350-2000 m/z at a resolution of 30 k (AGC target 1e6) and while the top 12 FT-HCD-MS/MS spectra were acquired in data dependent mode with an isolation window of 2.0 m/z and at a resolution of 15 k (AGC target 1e5) using a normalized collision energy of 30 and a 45 sec exclusion time.

*Growth assay of* Sulfitobacter *sp. N5S in co-culture with* P. tricornutum

*Sulfitobacter* sp. N5S (N5S) was isolated from Bodega Bay (Bodega Bay, California; 38.332615, -123.048296) by spreading whole seawater onto Marine Agar plates. Resultant colonies were picked, re-streaked and purified. A draft genome of the isolated N5S strain (NCBI Accession # PRJNA58103) was generated from DNA extracted with a DNAEasy kit (Qiagen, Germany) through the JGI's CSP (JGI Award doi: 10.46936/10.25585/60001054). Initial annotations and analyses were done through JGI's integrated microbial genomes (IMG) pipeline (64) and dbCAN2 (53) for carbohydrate-active enzyme annotation (version dbCAN12). The growth dynamics of N5S in co-culture with *P. tricornutum* was assessed by inoculating an axenic culture of *P. tricornutum* CCMP 2561 (CCMP 2561, National Center for Marine Algae and Microbiota; ncma.bigelow.org) with a sterile loop from a single colony. The co-culture was grown in batch mode and serially transferred for at least 5 transfers to establish an in-balance co-culture where N5S depends on *P. tricornutum* released organic matter (11). The co-culture was grown in borosilicate glass 13 mm diameter tubes on a 14/10 hr light/dark regime at 75 μmol quanta m-2 s−1 (cool white fluorescent) at 22 °C. The seawater (F/2) medium was prepared using Instant Ocean salts (35 g/L) with 880 μM nitrate (65).

Algal and bacterial abundance was assessed over a 10-day growth period on an Attune benchtop flow cytometer with a CytKick autosampler (Thermo Fisher Scientific, Waltham, MA) fitted with a blue laster (excitation 488 nm). One mL of each sample was collected and immediately fixed with glutaraldehyde (0.25% final concentration), flash frozen, and stored at −80 °C. Fixed samples were diluted in sterile-filtered media to achieve >4000 event counts per

second and stained with 1X SYBR Gold (Thermo Fisher Scientific, Waltham, MA) for 10 minutes in the dark prior to analysis. The instrument parameters were set up to threshold at 0.1 x 1000 on the BL1 detector, sample acquisition volume of 200 µL run at a flow rate of 100 µL/min. Gating was performed based on forward scatter, side scatter, and fluorescence of SYBR Gold (BL1 detector, emission/BP of 530/30). MilliQ blanks were run between treatment replicate sets to reduce carryover, and media blanks were included to correct for background particle noise.
Supplementary Figures

*Cross feeding experimental design*

To generate *P. tricornutum* lysate, a stock culture of axenic CCMP 2561 was inoculated into 125 mL sterile F/2 media (described above) in sterile, 200 mL acid-washed glass flasks with shaking (90 rpm, 22°C; 12 h : 12 h, light : dark; 3500 lux illumination). After 5 days of growth when the culture reached ~5,000 chlorophyll *a* relative fluorescence (RFU, measured using the BioTek Cytation 5 plate reader [Agilent]), or early-log growth (19), algal cells were pelleted by spinning at 5000 rpm for 8 minutes and the spent media was discarded to remove algal exudate. The cells were resuspended in fresh F/2 media. Then, the cells in fresh media were lysed using an Ultrasonic Processer XL sonicator (Misonix) using the following protocol: sonicate 3x at level 3 for 30 s, then 3x at level 5 for 30s. The lysate was kept on ice in between each round for 1-2 minutes to allow the sample to cool. Finally, the lysate was filtered through a 0.8 µm syringe filter to remove large intact cells and debris.

The final lysate was then incubated with or without OT1 to produce “abiotic” and “OT1 conditioned” lysate, respectively. Here, OT1 cells grown to exponential phase in Zobell marine broth (63) were pelleted by spinning at 5000 rpm for 8 minutes, spent media thoroughly removed, and resuspended in fresh F/2. The conditioning experiment was done in triplicate per condition (OT1 conditioned *versus* abiotic) in 3 mL sterile borosilicate tubes at 22°C in the dark. OT1 conditioned lysate was generated by inoculating the lysate to an OD600 of ~0.12. Abiotic lysate was generated by adding nothing to the lysate. Zobell broth was used as a positive control. Growth was tracked using OD600 with the BioTek Cytation 5 plate reader (Aglient) and samples for flow cytometry were collected and processed as described in Supplemental methods.

A kinetic curve was generated for OT1 and N5S grown in the abiotic and OT1 conditioned lysate. Here, the abiotic/conditioned lysates were filtered through a 0.2 μm syringe filter. Exponentially growing OT1 or N5S in Zobell marine broth was pelleted by spinning at 5000 rpm for 8 min, and then the spent media was thoroughly removed. Then, each isolate was resuspended in either abiotic lysate, OT1 conditioned lysate, or Zobell marine broth and aliquoted into 5 replicate wells per condition in a 96 well plate (rounded bottom). A negative control for each treatment was run. OD600 was read every 20 min for 72 h using the BioTek Cytation 5 plate reader (Agilent) at 22°C in the dark. Samples for flow cytometry were collected at T=0 h and T=72 h in triplicate and processed as described in the Supplemental Methods. The t-test was used to compare cell abundances at T= 72h within abiotic *versus* OT1 conditioned lysate using GraphPad Prism 9 and data were plotted using ggplot2 (42).

### Supplementary Figures

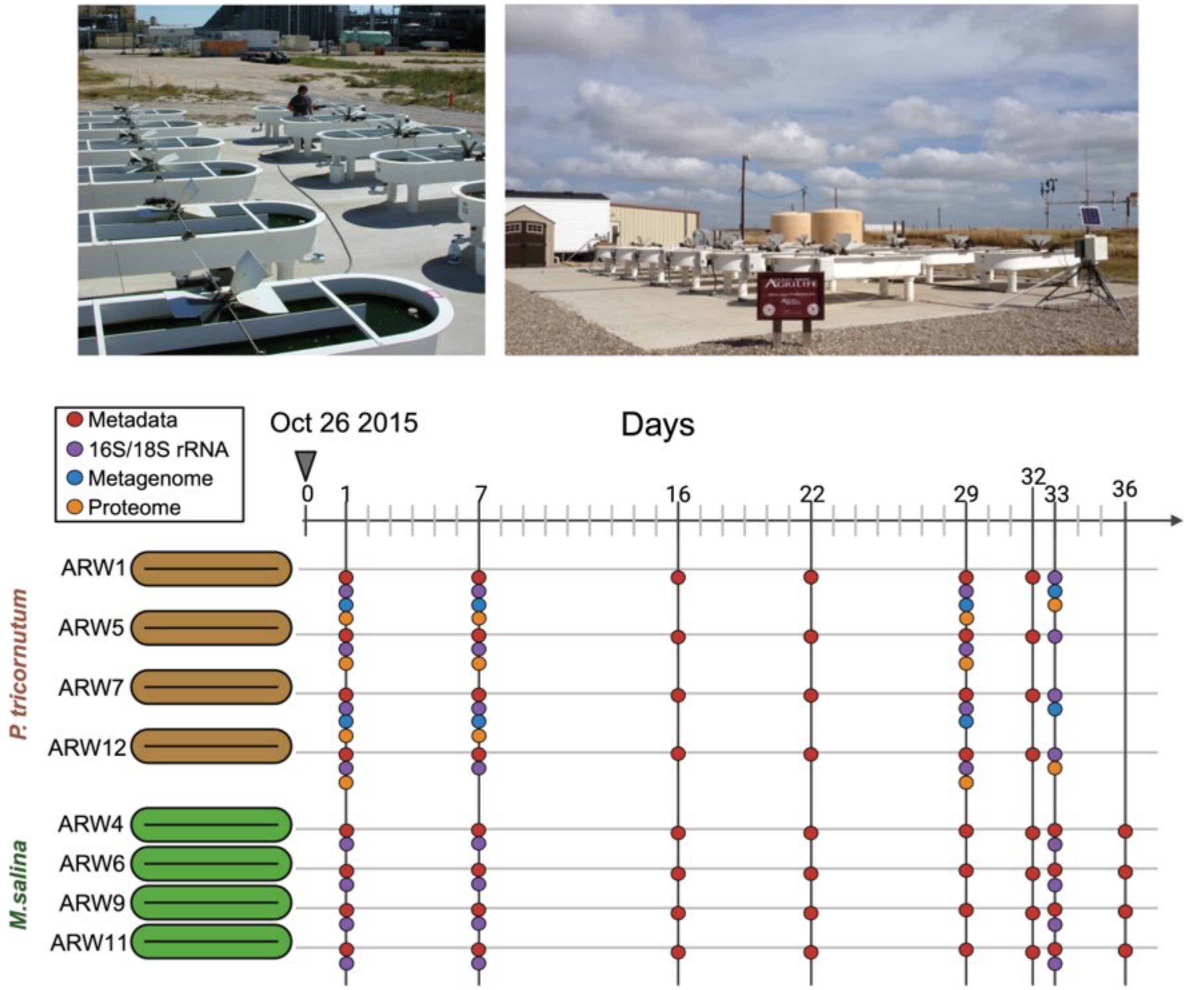


**Supplementary Figure 1.** (top) The Algal Raceway (ARW) Ponds housed at Texas A&M used for the study. (bottom) Schematic illustrating sampling timepoints for *P. tricornutum* ARW ponds, *M. salina* ARW ponds. The ponds were set up in four replicates for each host. The samples collected for each pond and day are shown as color-coded circles. All metadata measurements ended for *P. tricornutum* on Day 32 due to visual clearing of all four ponds. The arrow shows the day ("Day 0") of inoculation for the ponds. The image was designed using Biorender (biorender.com).

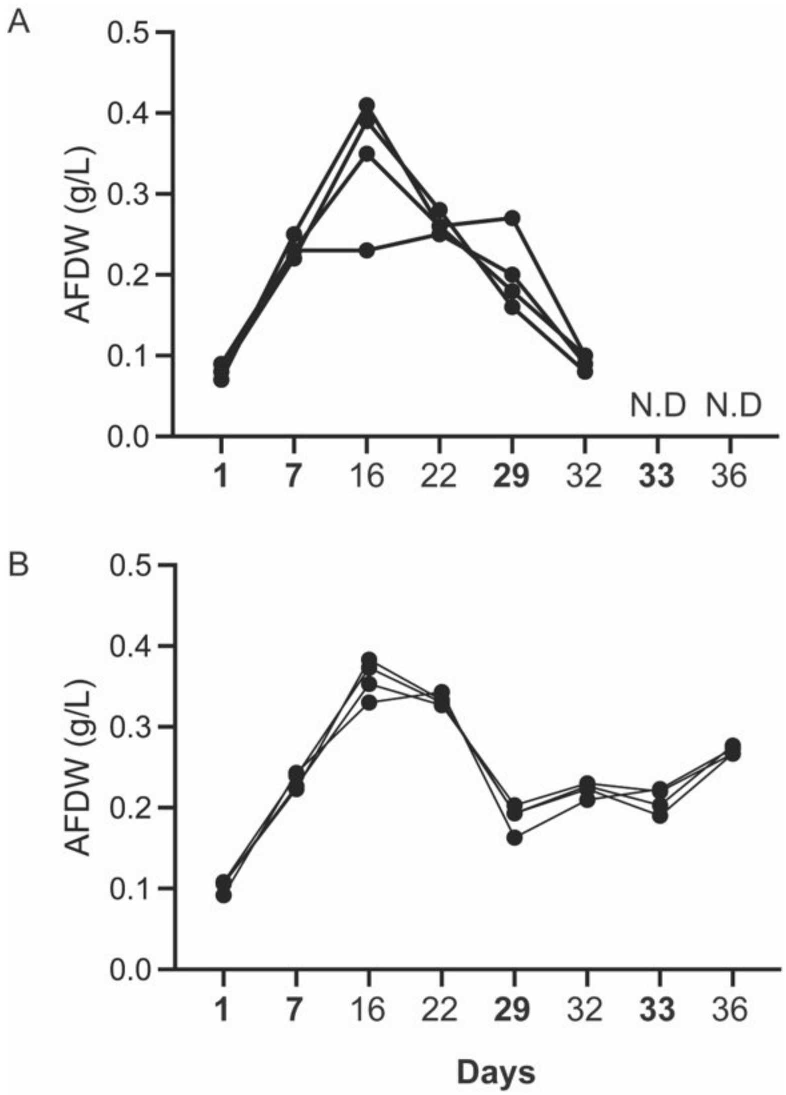


**Supplementary Figure 2.** Biomass measurements of the A) *P. tricornutum* and B) *M. salina* ponds. Replicate pond measurements are shown as separate lines. No further biomass measurements were conducted for Day 33 and 37 ("N.D") for *P. tricornutum* due to the raceways having visibly cleared. Dates emboldened indicate sampling timepoints for sequencing data.

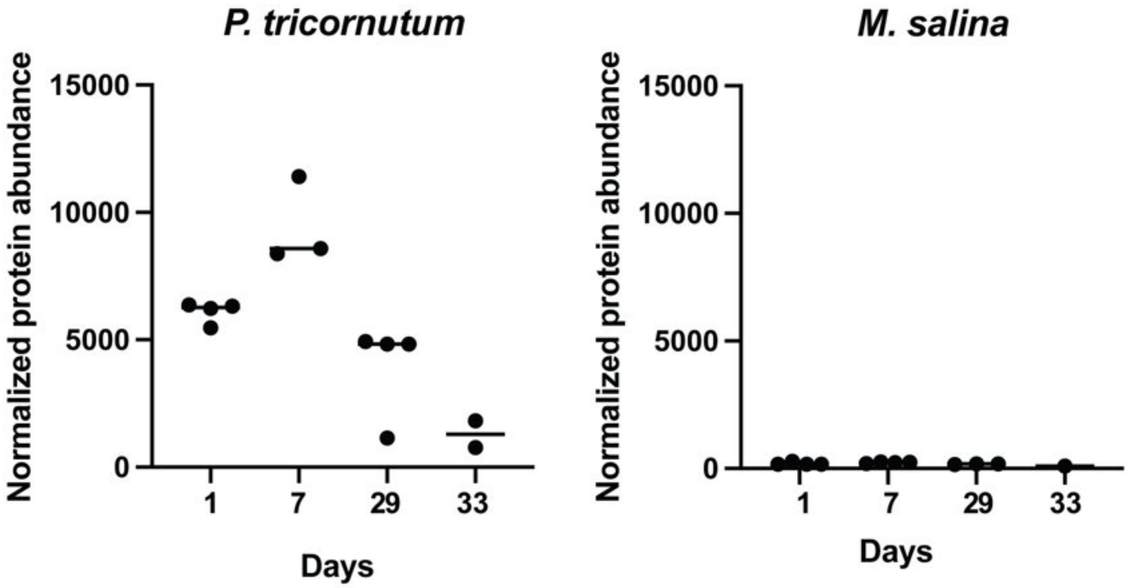


**Supplementary Figure 3.** Total protein abundance, derived from *P. tricornutum* inoculated pond metaproteomes, mapped to *P. tricornutum* (left) *versus M. salina* (right) genome.

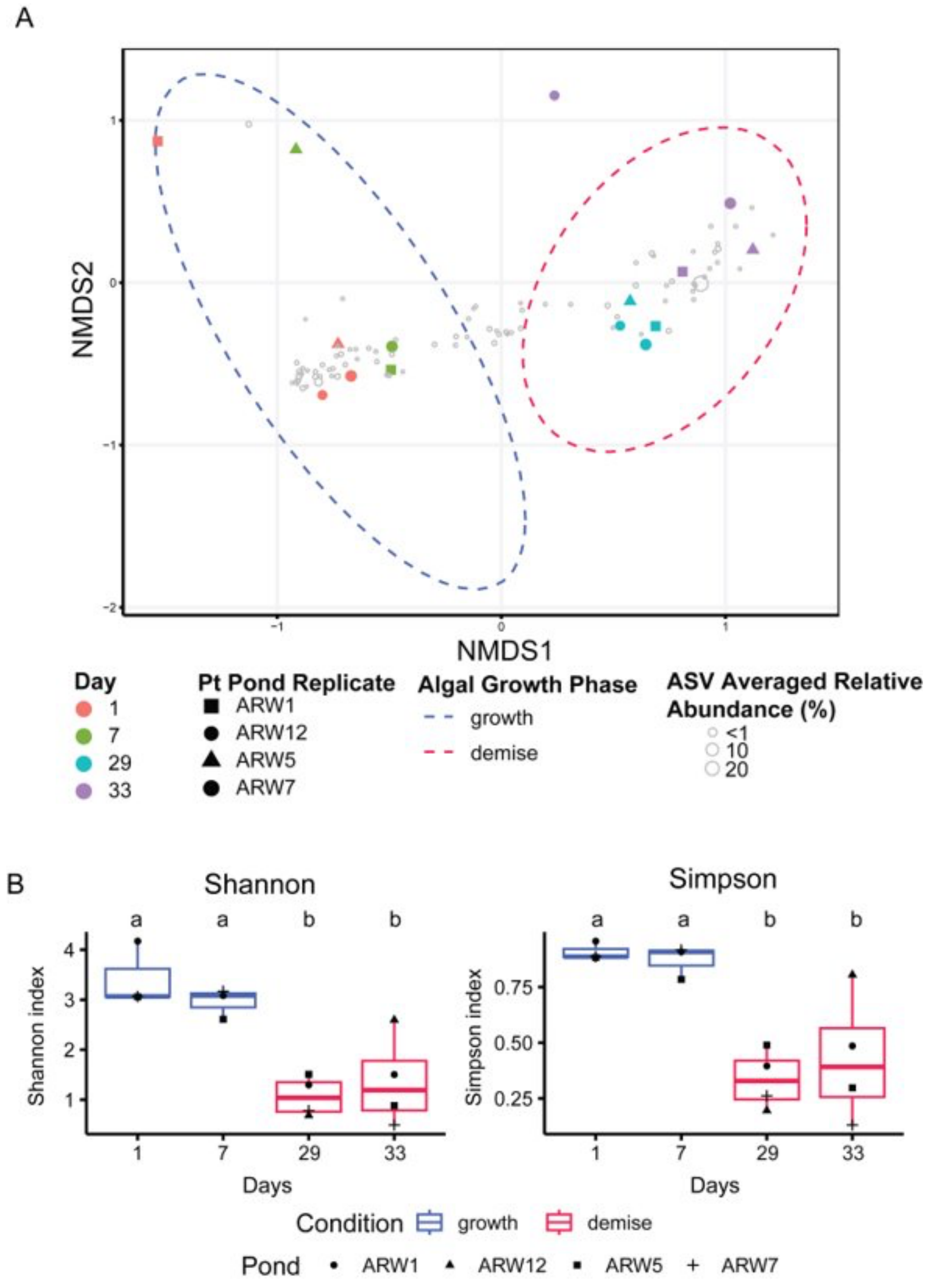


**Supplementary Figure 4*. P. tricornutum* Pond 16S rRNA community structure.** A) NMDS ordination based on the Bray-Curtis dissimilarity matrix of 16S rRNA bacterial communities across the *P. tricornutum* time course. The 95% confidence ellipses (stat_ellipse, Fox and Weisberg 2011) are grouped by algal condition (Growth vs Demise, ADONIS p<0.005). ASV's are plotted as transparent bubbles based on their NMDS ordination coordinates and are scaled based on their averaged relative abundances across all samples. B) Alpha diversity results of the *P. tricornutum* microbiomes, summarized by timepoint and color coded by algal growth phase. Letters indicate statistical differences based on Tukey's HSD adjusted p-values.

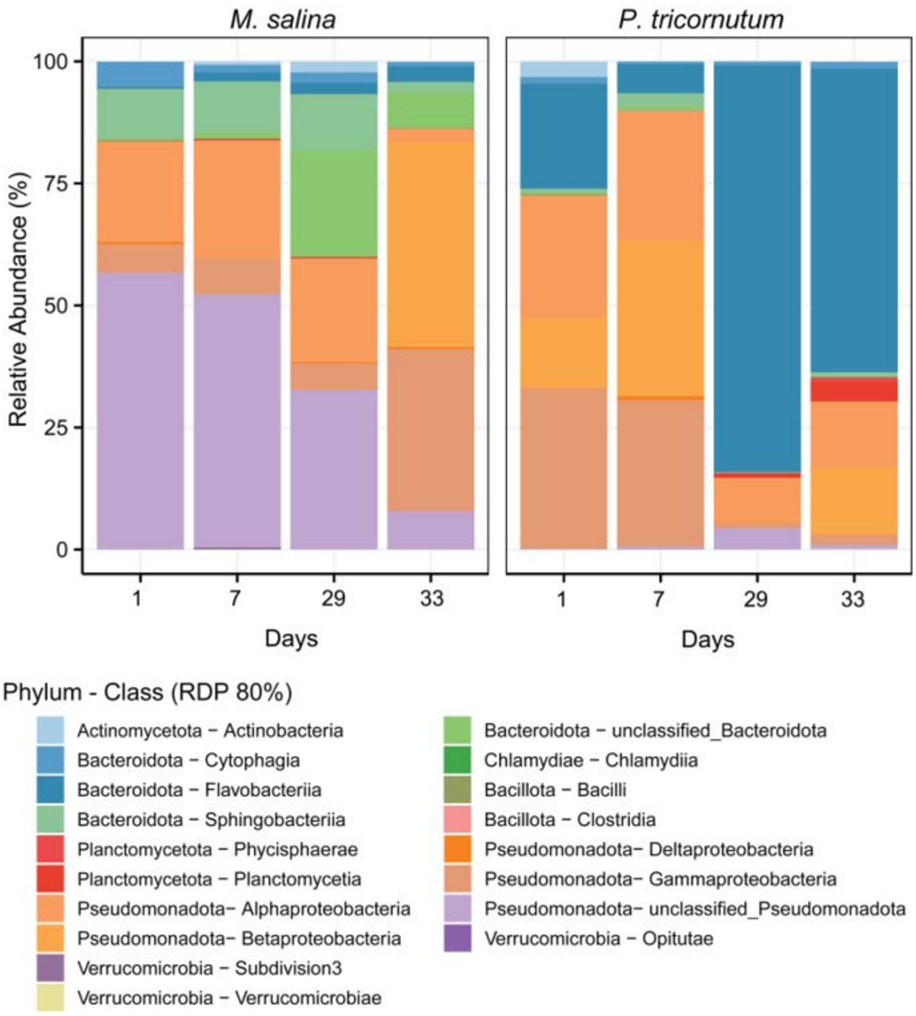


**Supplementary Figure 5.** Bacterial community structure of ponds inoculated with either *M. salina* or *P. tricornutum,* shown as the relative abundance of 16S rRNA ASV's grouped by Class and summarized by Day samples (n=4/day)

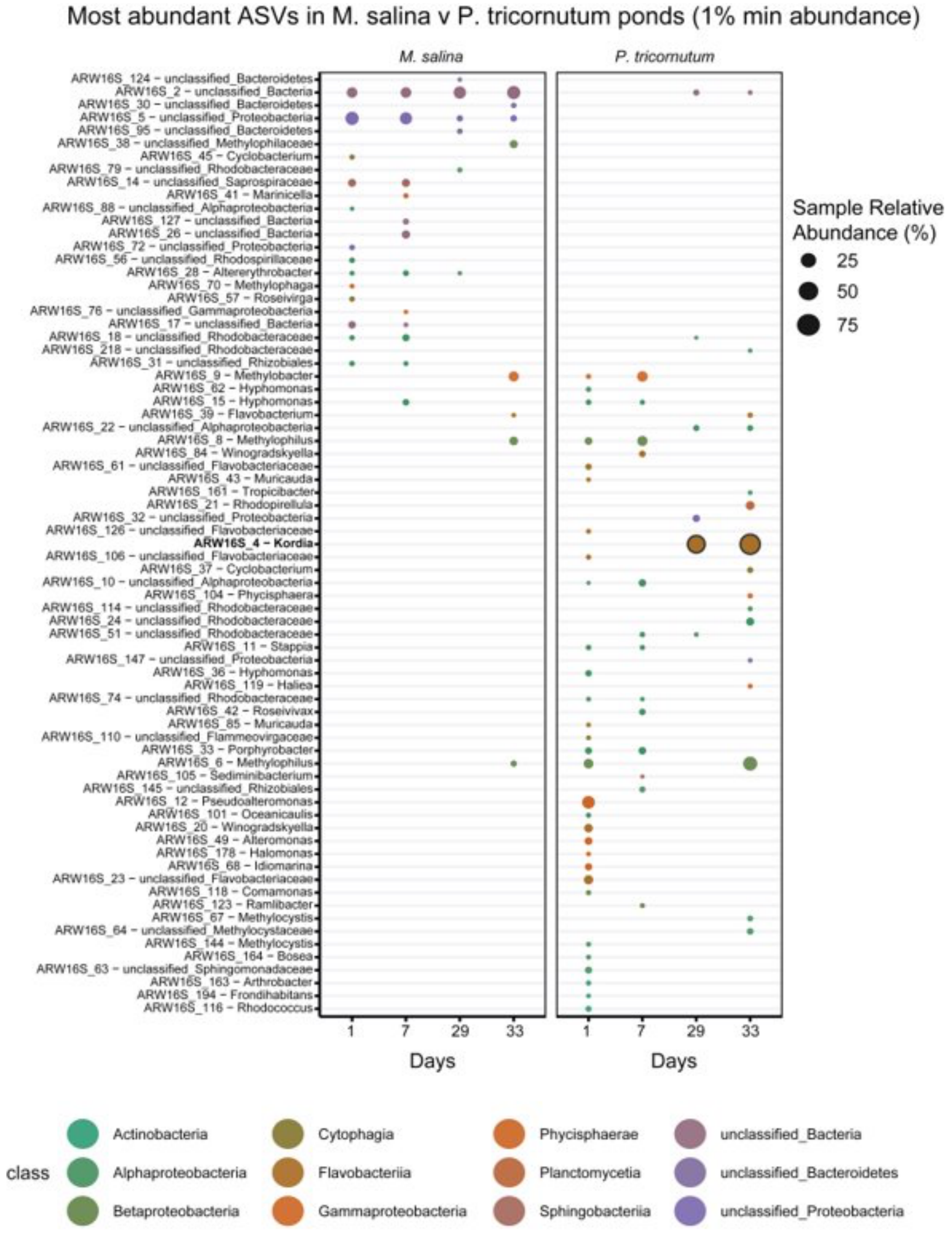


**Supplementary Figure 6.** Comparison of ASV-level relative abundance between *M. salina* vs *P. tricornutum* inoculated ponds. Sample relative abundance is averaged by timepoint (n=4). ASVs with a minimum of 1% averaged relative abundance value are shown. Bubbles assigned to *Kordia* (ARW16S_4) are emboldened for visual purposes.

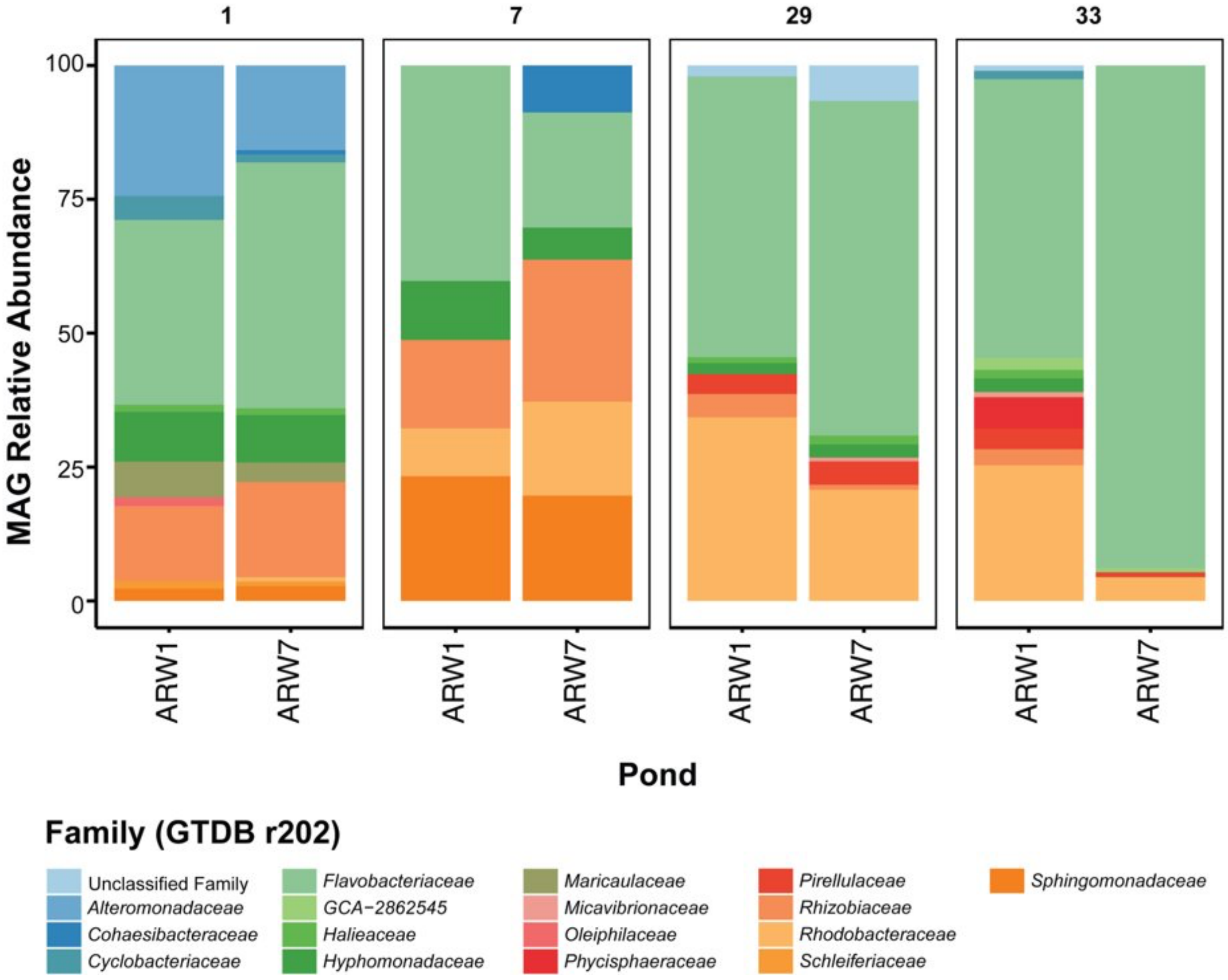


**Supplementary Figure 7.** Abundance of assembled MAGs across metagenomes, shown by family-level proportions and grouped by time. Both replicate ponds sequenced for metagenomes are shown. Abundance of MAGs was calculated using the mean of the median fold coverage (see Methods) and are plotted as relative abundances.

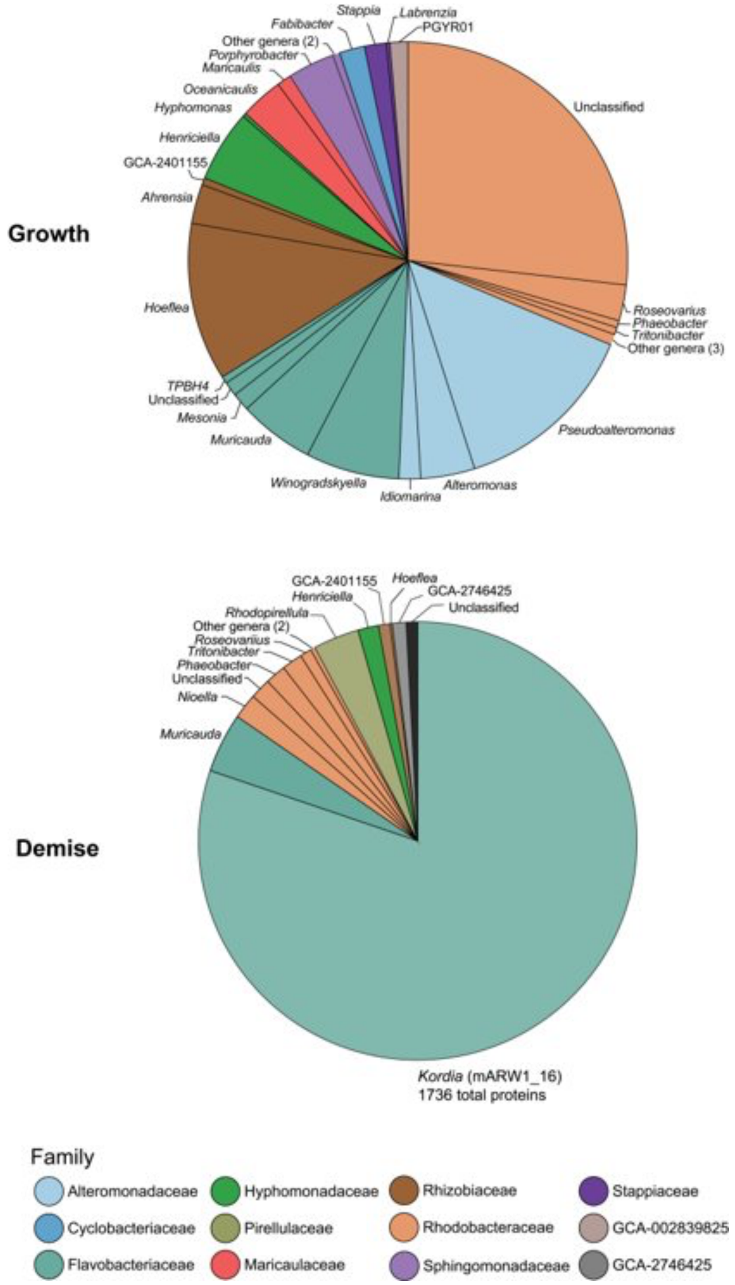


**Supplementary Figure 8.** Taxonomic breakdown of the number of detected proteins in the Growth (days 1 and 7) versus Demise (days 29 and 33) phase. A protein was considered "detected" if the averaged NSAF was >= 2 within each algal growth condition samples. Charts are color coded by Family-level taxonomic assignment and grouped by Genus.

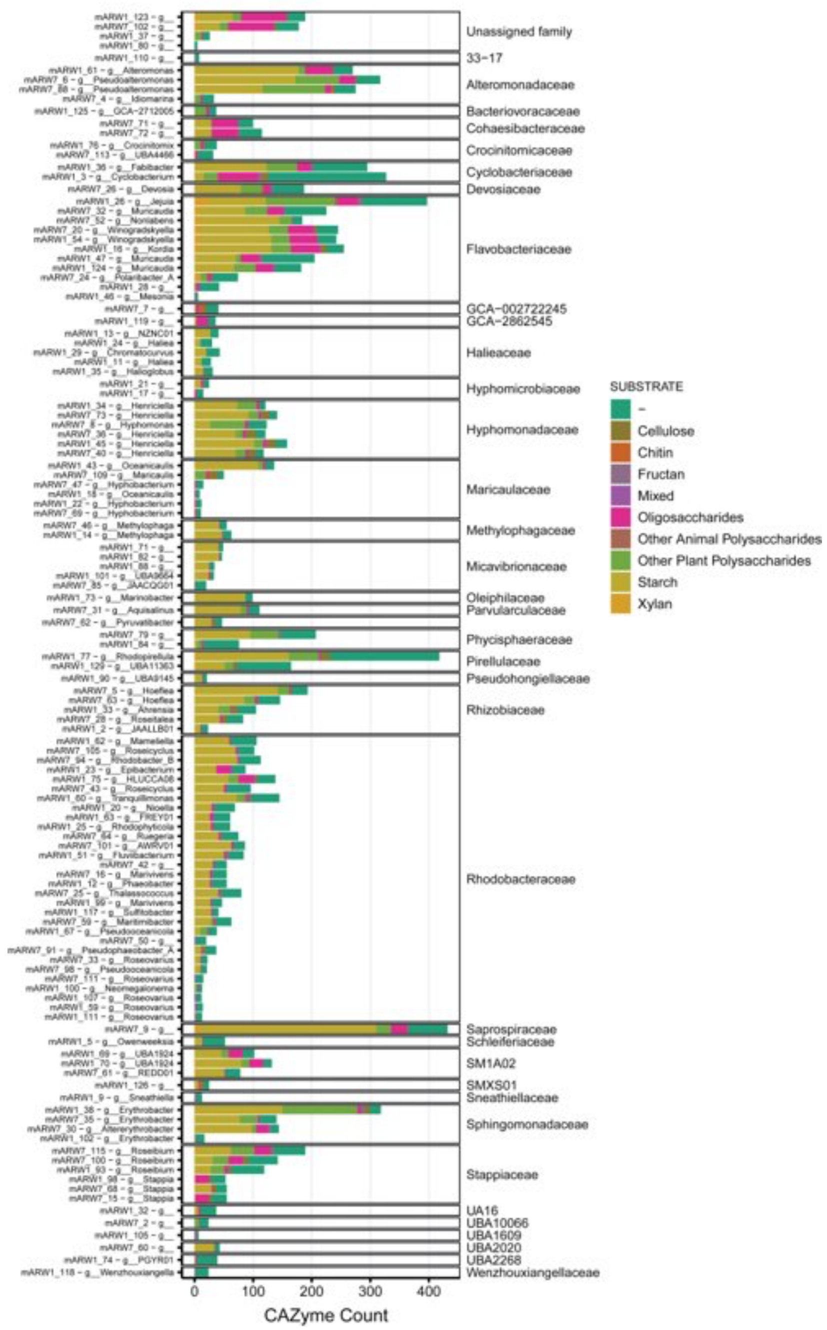


**Supplementary Figure 9.** Distribution of CAZymes (Glycoside hydrolases and polysaccharide lysases) across all ARW1 and ARW7 assembled MAGs, grouped by Family and color coded by predicted substrate.

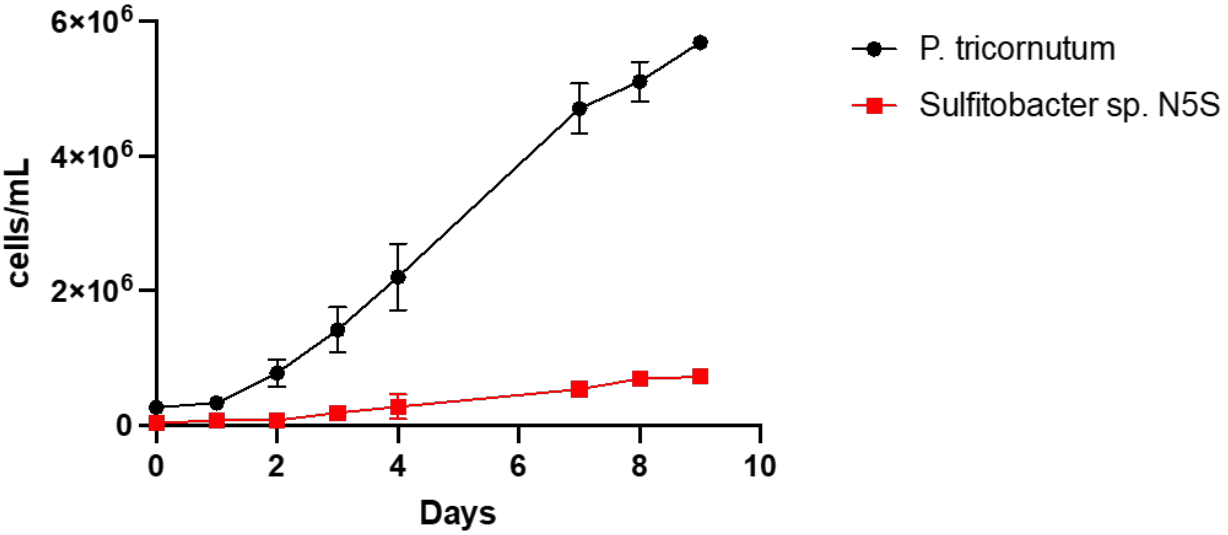


**Supplementary Figure 10.** Growth of *P. tricornutum and  Sulfitobacter* sp. N5S in co-culture. Error bars represent the standard deviation.

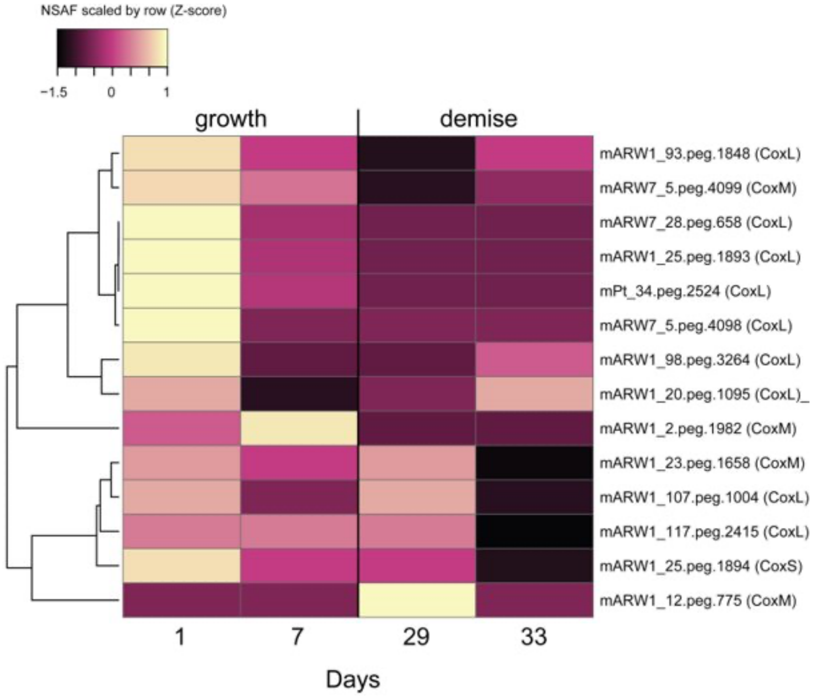


**Supplementary Figure 11.** Expressed carbon monoxide oxidation proteins in the *P. tricornutum* ponds along the time course. Proteins with >= 3 total counts are shown. Normalized protein abundances (nsaf) were averaged by timepoint and are shown as relative values (z-score) across timepoint. Proteins are clustered by Average Linkage and distances measured by Pearson correlation.

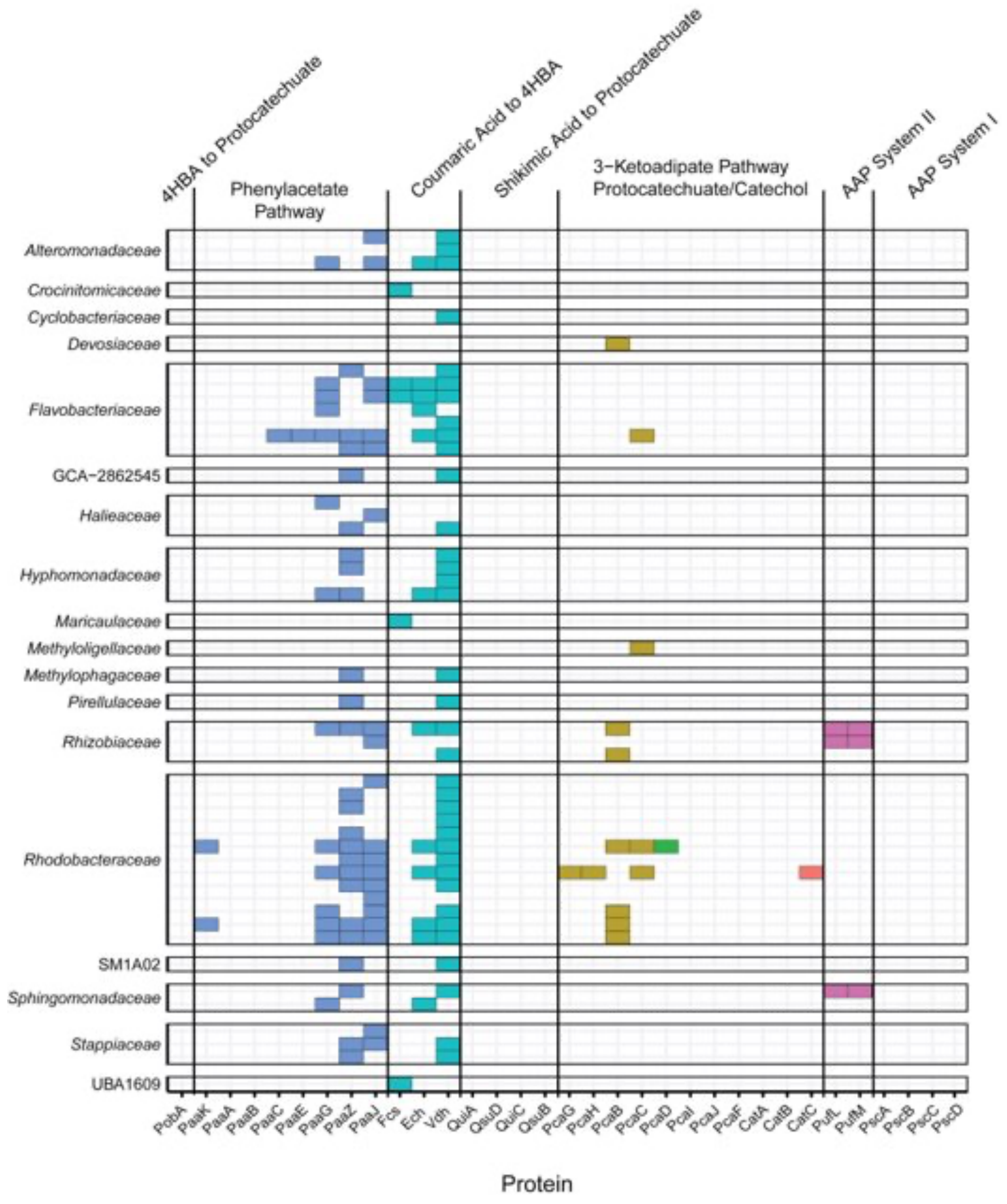


**Supplementary Figure 12.** Detection of expressed proteins for other pathways of involved in alternative carbon and energy metabolism (Aromatic compound degradation [Phenylacetate pathway, coumaric acid to 4HBA, Shikimic acid to protocatechuate, 3-Ketoadipate pathway] and Light [AAP System I and II]).
